# DNA methylation calling tools for Oxford Nanopore sequencing: a survey and human epigenome-wide evaluation

**DOI:** 10.1101/2021.05.05.442849

**Authors:** Yang Liu, Wojciech Rosikiewicz, Ziwei Pan, Nathaniel Jillette, Ping Wang, Aziz Taghbalout, Jonathan Foox, Christopher Mason, Martin Carroll, Albert Cheng, Sheng Li

## Abstract

**Background:** Nanopore long-read sequencing technology greatly expands the capacity of long-range single-molecule DNA-modification detection. A growing number of analytical tools have been actively developed to detect DNA methylation from Nanopore sequencing reads. Here, we examine the performance of different methylation calling tools to provide a systematic evaluation to guide practitioners for human epigenome-wide research.

**Results:** We compare five analytic frameworks for detecting DNA modification from Nanopore long-read sequencing data. We evaluate the association between genomic context, CpG methylation-detection accuracy, CpG sites coverage, and running time using Nanopore sequencing data from natural human DNA. Furthermore, we provide an online DNA methylation database (https://nanome.jax.org) with which to display genomic regions that exhibit differences in DNA-modification detection power among different methylation calling algorithms for nanopore sequencing data.

**Conclusions:** Our study is the first benchmark of computational methods for mammalian whole genome DNA-modification detection in Nanopore sequencing. We provide a broad foundation for cross-platform standardization, and an evaluation of analytical tools designed for genome-scale modified-base detection using Nanopore sequencing.

## Background

DNA methylation, the process by which methyl groups are added to DNA molecules, is a fundamental epigenetic modification process in gene transcription regulation [1]. Several DNA modifications, such as N6-methyladenine (6mA), N4-methylcytosine (4mC), and 5-methylcytosine (5mC) and its oxidative derivatives, are diversely distributed in genomes and play important roles in genomic imprinting, chromatin structure modulation, transposon inactivation, stem cell pluripotency and differentiation, inflammation, and transcription repression regulation [2–4]. DNA methylation measurement has traditionally depended on the combination of bisulfite conversion (which can damage DNA) and next-generation sequencing, which only detects short-range methylation pattern [5].

Recently, third-generation sequencing technologies, including single molecule real-time (SMRT) sequencing by Pacific Biosciences (PacBio), and Nanopore sequencing by Oxford Nanopore Technologies (ONT), have overcome the length limitation to achieve ultra-long read, single-base detection at a genome-wide level [6, 7]. SMRT sequencing can detect 5mC based on polymerase kinetics at 250x coverage [8]. This is due to the subtle impact of 5mC on polymerase kinetics [8]. Thus, the high coverage requirement and direct single-molecule 5mC detection by SMRT is still challenging [9]. Single molecule real-time bisulfite sequencing allows to sequence up to ∼2kb amplicons but it relies on bisulfite conversion [10].

Nanopore sequencing, instead of using a sequencing-by-synthesis method to detect signal for the amplified DNA fragment population, is able to directly detect DNA or RNA translocation through a voltage-biased Nanopore sensor, enabling rapid long-read sequencing and single-base and single-molecule sensitivity [11]. Several different versions of Nanopore chemistry have been developed by ONT to improve the accuracy of single-cell molecular identification (**Figure 1A**). The initial pore version of flow cells, termed R6/R7, was replaced by R9 pore series. R9 pore series were derived from the bacterial amyloid secretion pore gene Curlin sigma S-dependent growth (CsgG) to yield a modal (i.e., most commonly observed) accuracy of up to 95% at the single-molecule level [12, 13]. Q scores, also known as Phred quality scores, are logarithmically linked to the error probability (P) of each called base: *Q* = −10 × *log*_10_(*p*). Q scores measure the accuracy of nucleobase identification in DNA sequencing. Higher Q values correspond to lower error probability and higher quality [14, 15]. For example, Q30 indicates that the chance that a specific base is called incorrectly is 1 in 1000. R9-series pores, R9.4 and its slightly updated, broadly used version R9.4.1, are the most favored version and can achieve the best consensus accuracy at 99.99% (Q45) [16, 17]. Recently, ONT released Nanopore R10 with a predicted model accuracy of 94% [18, 19], and introduced the newest version R10.3 with of 99.995% single molecule consensus accuracy, which has a longer barrel and a dual-reader head inside the pore [15, 17]. The current study is conducted on R9.4 series version.

**Figure 1.**
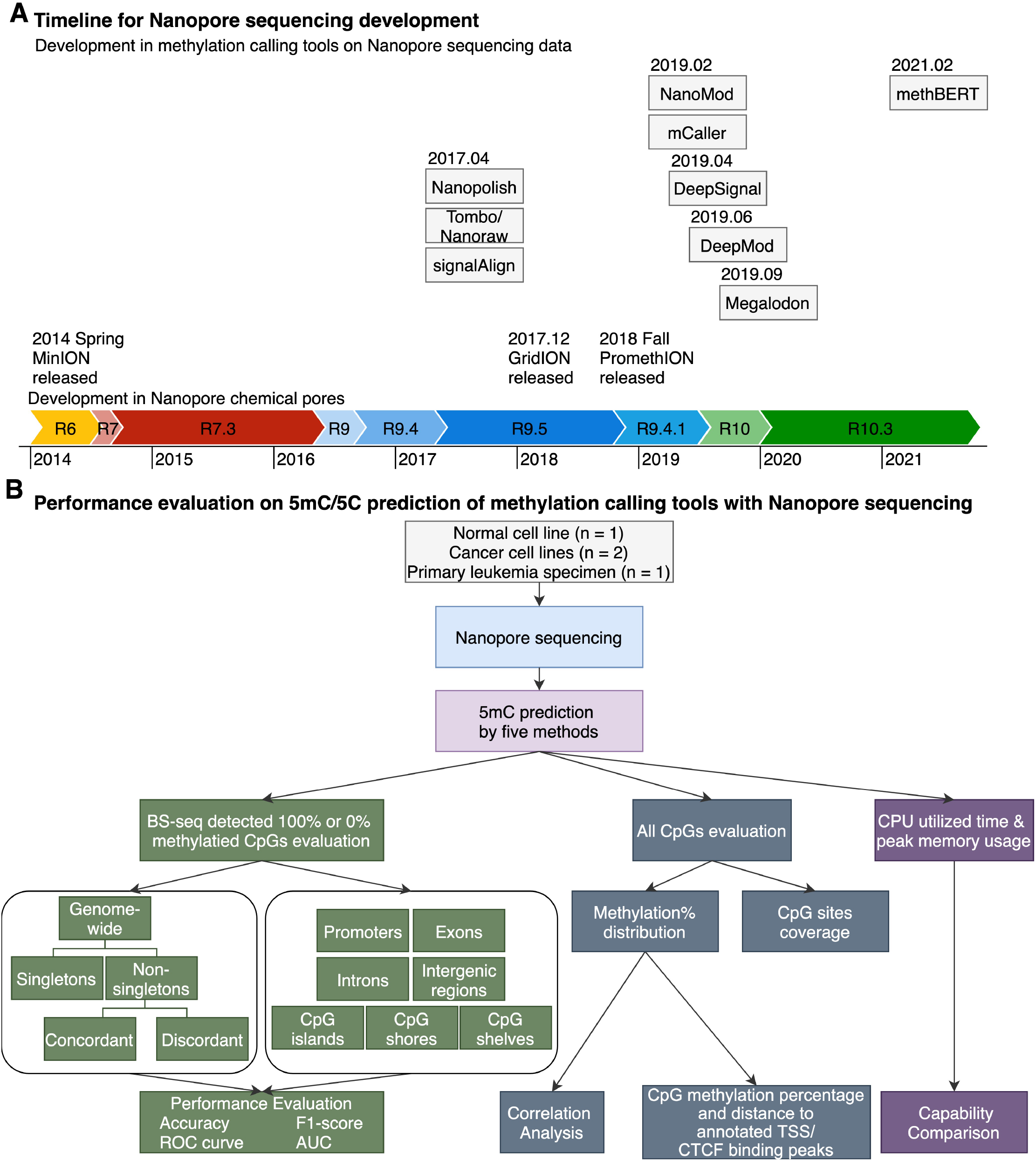
Technological development of methylation calling tools and performance evaluation flowchart. (**A**) Timeline of publication and technological developments of Oxford Nanopore Technologies (ONT) methylation calling tools to detect DNA cytosine modifications. Methylation calling tools are listed acc in order by publication date instead of bioRxiv online submission date (BioRxiv date for methBERT and Github release time for Megalodon since they lack an available official publication). Chemical pore versions of Nanopore flow cell are represented as colored bars. Chemical pore version of Nanopore flow cell compatibility for each methylation calling tools is shown in corresponding colors. Relevant publication time are from multiple source [9, 15, 17, 18, 21, 49–51, 78, 88, 92–94]. (**B**) Performance evaluation on 5mC/5C prediction of methylation calling tools with Nanopore sequencing. We generated four datasets for nanopore sequencing, applied five methylations calling tools separately to detect methylation status and compare the 5mC/5C classification with the background truth BS-seq. To compare the performance of different tools, we compared methylation calling results in singletons/non-singletons regions or biological relevant genomic regions, correlated all CpGs sites with methylation distribution in BS-seq, and evaluated the running speed and computing memory usage.

Nanopore sequencing techniques enables DNA modification detection due to the difference in the electric current intensity produced from a nanopore read, termed “squiggles”. Specifically, the ionic-current resulting from the passage of modified bases through the pores differs from the current produced by the passage of unmodified bases [14, 20]. The difference can be determined after nanopore read base calling and alignment by: (1) statistical tests comparing to an in silico reference or a non-modified control sample [21, 22]; (2) pre-trained supervised learning models such as neural network [23–25] and Hidden Markov Model (HMM) [9, 26]. However, DNA-methylation detection using Nanopore data presents a methodological challenge, i.e., accurate detection of DNA modifications in CpG sites (CpGs) termed non-singletons. A 10-base-pair (bp) region that contains only one CpG site is defined as a singleton, while a 10-bp region that contains more than one CpG site are called non-singletons [9]. The primary difficulty is the capacity to detect modifications in different CpGs that are in close proximity to one another on a DNA fragment, as it is assumed that all CpGs within a 10-bp region share the same methylation status. Several methylation calling tools have been developed to handle singletons to improve DNA-methylation detection accuracy (**Table 1**), but DNA-methylation detection power for non-singletons containing both methylated and unmethylated states remains difficult [9, 27]. Also, DNA methylation level is not linearly distributed across the genome and is dependent on genomic context [28–30]. Therefore, the accuracy of methylation callers likely differ among various types of genomic regions within which CpGs are located. However, there is no published guideline and systematic comparison of current DNA methylation calling tools for Nanopore sequencing using human natural DNA [31], especially at whole epigenome scale [32, 33].

**Table 1.**
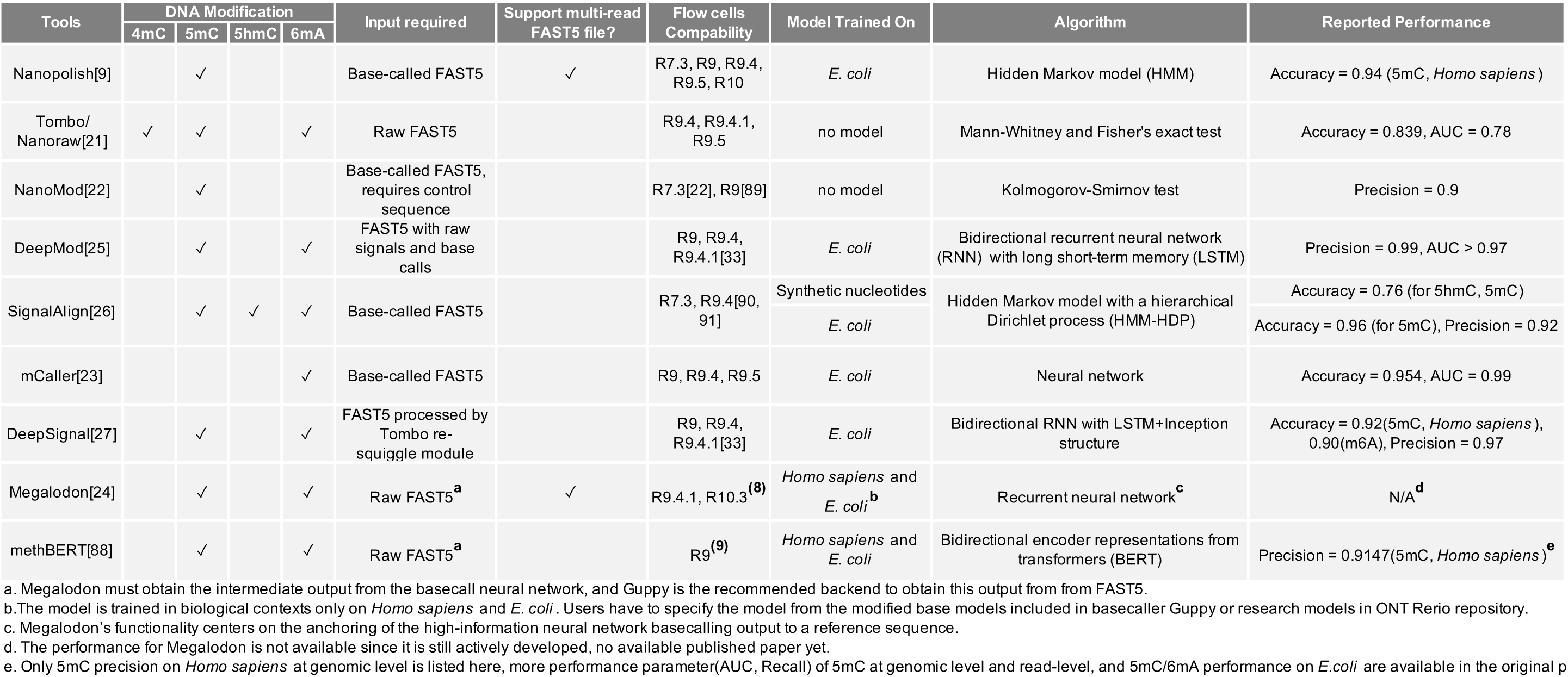
Current DNA methylation-calling tools for Nanopore sequencing. Reference is shown in the table correspondingly[9, 21–27, 33, 88–91].

Here, we present the first benchmark of computational methods for detecting of DNA 5-methylcytosine (5mC) from human Nanopore sequencing data at whole genome scale. We assess the impact of CpG locations on detection accuracy using human whole genome nanopore sequencing data, with a focus on the impact of genomic context and singletons vs. non-singletons. It has been reported that even homogeneous cell populations can exhibit cell-to-cell variations in epigenetic pattern (epiallele) such as gain or loss of cytosine methylation at specific loci [34]. Such epigenetic heterogeneity is increasingly recognized as a contributor to biological variability in tumors and worse clinical outcomes in malignancies [5]. Thus, to enable assessment of this critical epigenetic heterogeneity, we have evaluated the DNA methylation accuracy at single-molecule and single-base resolution, which is critical for epigenetic heterogeneity assessment [35–38]. This comprehensive survey and systematic comparison offer user-specific, best-practice recommendations to maximize accurate detection of 5mC using current methylation calling tools and provide guidance for next generation calling tools. We also generated and made available a R Shiny database to distribute the modification-detection power associated with different genomic regions using different tools for development of future algorithms and analytic tools development.

## Results

### Benchmarking dataset

We used four datasets for benchmarking: Nanopore sequencing of the human B-lymphocyte cell line NA19240 (hereafter referred to as NA19240 in the following text) [39], human leukemia cell lines K562 and HL-60 (referred to as K562 and HL-60), and a human primary acute promyelocytic leukemia clinical specimen (referred to as APL).

NA19240 was sequenced at ∼32x coverage by the 1000 Genomes Project [39] as a high-coverage dataset. We take the union of sites from two reduced representation bisulfite sequencing (RRBS) replicates for NA19240 as corresponding DNA methylation ground truth. We generated nanopore sequencing data for K562, HL-60, and APL with ∼1-3x coverage and whole genome bisulfite sequencing (WGBS) for APL. We used the published WGBS for K562 and RRBS for HL-60 as ground truth.

### Overall strategy to compare DNA methylation calling tools

Several methylation calling tools have been developed to detect DNA methylation using Nanopore direct DNA sequencing data (**Table 1**). Among the nine tools, seven tools are compatible with R9.4 series flow cells and six of these tools can predict 5-methylcytosine (5mC). To compare the performance of these state-of-the-art methylation calling tools, we developed a three-step standardized workflow to compare five methylation calling tools targeting 5mC in CpG context compatible with the most favored Nanopore flow cell version (R9.4.1 pores): Nanopolish [9], Megalodon [24], DeepSignal [27], Tombo/Nanoraw (referred to as Tombo) [21], and DeepMod [25] (**Figure 1B, Figure S1**). Nanopolish, Megalodon, DeepSignal and DeepMod, is model-based while Tombo is statistics-based. We excluded SignalAlign [26], as its repository has been no longer updated for over four years.

#### Step 1. Base-calling and quality control

To translate raw signal data into nucleotide sequences, we conducted the base calling step for Nanopore reads with Guppy (v4.2.2). Then we used NanoPack [40] for data visualization and processing, in order to assess the read length and quality of the base-called and to demultiplex sequencing data for downstream analysis. The APL, K562, and HL-60 ONT datasets exhibited comparable read length and base quality compared to the published NA19240 ONT dataset [39] (**Figure 2A-B**). Distribution of CpG sites distribution based on singletons/non-singletons is shown in (**Table S1, Figure S2**), while the number of CpG sites in various genomic contexts distribution is shown in **Figure 2C** and **2D**.

**Figure 2.**
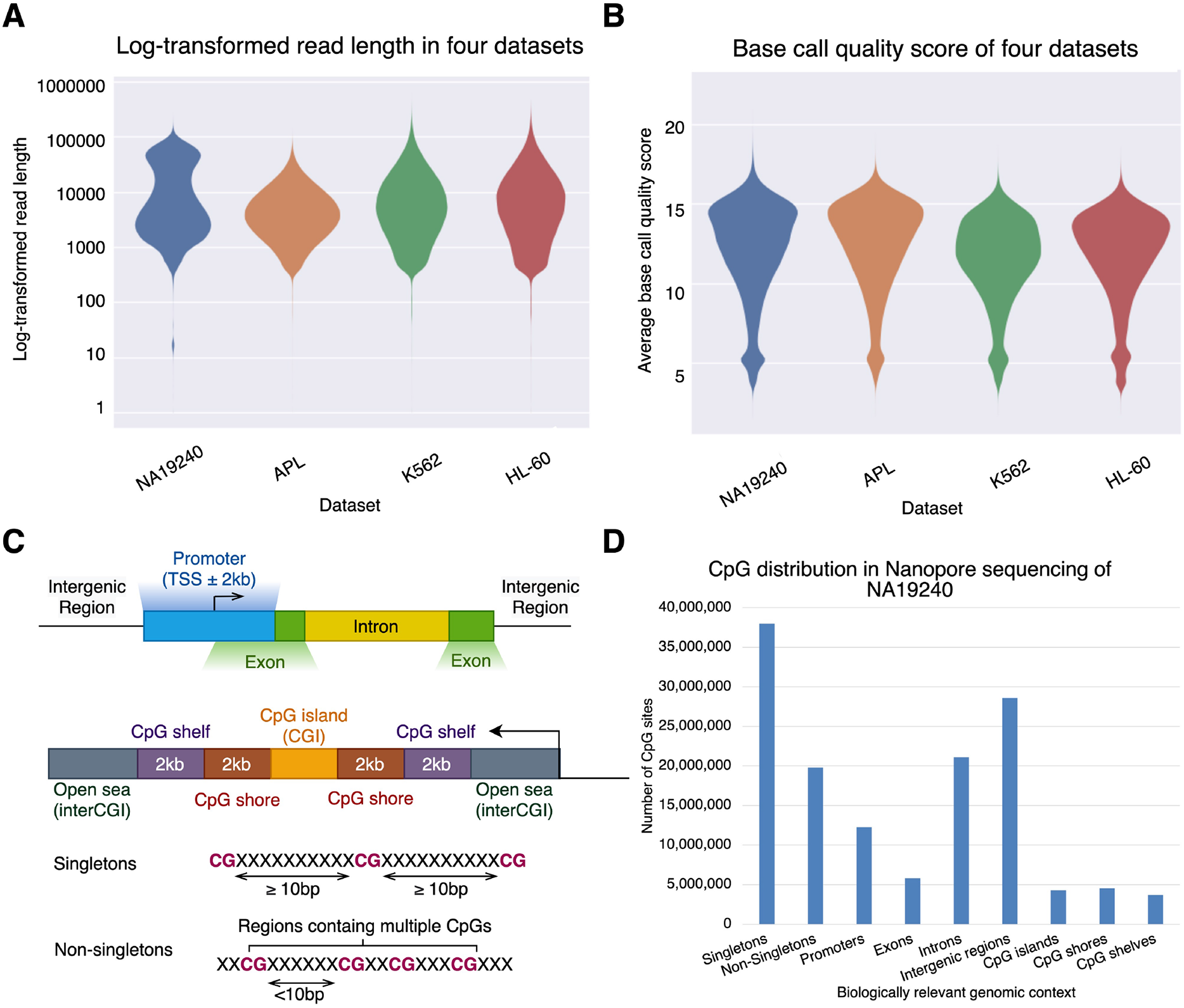
Datasets description. (**A-B**) Quality control summary for four datasets. (**A**) Violin plot of log-transformed read length (**B**) Violin plot of base call quality. Data shown are colored by dataset and plotted by Plots NanoPack [40]. **(C)** Scheme for biologically relevant annotations of genomic context and singleton/non-singleton classification. We consider biologically relevant genomic context including promoter, exon, intron, intergenic regions and CpG island. Singletons are CpG sites that contain only one CG within the 10 base pairs, including absolute state (CpG is 0% or 100% methylated) or mixed state (0% < CpG methylation frequency < 100%). Non-singletons are those CpGs that contain more than one CpG in 10bp. **(D)** CpG sites distribution based on singletons, non-singletons and biologically relevant genomic context in Nanopore sequencing of the NA19240 dataset. Only regions with coverage >= 3 were considered.

#### Step 2. Genome assembly and polishing

We aligned the base-called reads to the human genome assembly GRCh38/hg38 using minimap2 [41] for all five tools. The electric current signal level data of a nanopore read produced by an ONT sequencer is called a squiggle. Base calling a squiggle, i.e., translating the current signal into a DNA sequence, typically contains some errors when comparing to a reference sequence [42]. The Tombo re-squiggle algorithm refines the assignment from a squiggle to a reference sequence after base-calling and alignment. The re-squiggle algorithm is required by Tombo and DeepSignal for DNA methylation calling.

#### Step 3. Methylation calling

We detected 5mCs in CpG context with five methylation calling tools: Nanopolish [9], Megalodon [24], DeepSignal [27], Tombo [21], and DeepMod [25]. We then designed three performance evaluation criteria (**Figure 1B and S1**) to compare the performances of each methylation calling tools. First, we evaluated the predictions of 5mCs at single-molecule, single-base resolution based on per-read prediction accuracy of fully methylated or fully unmethylated CpG sites determined by bisulfite sequencing data. We examined various biologically relevant genomic regions and singletons vs. non-singletons. Singletons are CpG sites with only one CpG within the 10-bp region, while non-singletons contain more than one CpG with the 10-bp region [32]. We further divided non-singletons into two sub-categories: (1) concordant non-singletons: all CpGs within the region share the same absolute methylation state (i.e., all fully methylated or all fully unmethylated), (2) discordant non-singletons: the methylation states of CpGs appearing in a close neighborhood (10bp) were mixed with both fully methylated and fully unmethylated sites present. Second, we measured the 5mC methylation correlation coefficient between ONT output and bisulfite sequencing data across all CpG sites at genome level. Third, we assessed the running speed and per-read resource usage evaluation. Further details on performance criteria used in evaluation are shown in **Methods**.

### Predictions of 5mC at single-molecule, single-base resolution

To understand the impact of various DNA methylation callers on 5mC prediction at single-molecule, single-base resolution, from different genomic contexts, we assessed the per-read accuracy in singletons and non-singletons. We compared methylation-calling performances on fully methylated/unmethylated CpGs in bisulfite sequencing (BS-seq) (coverage>=5) at the singleton and non-singleton levels across four datasets (**Table S2**). For NA19240, there are 30,377 singleton CpGs and 224,645 non-singleton CpGs in BS-seq (coverage>=5) that overlap with the ONT data. The comparison performance metrics include accuracy, F1 score, receiver operating characteristic curves (ROC curves) and area under the ROC curve (AUC) (**Figure 3, Figure S3, and Table S3**). DeepMod performance is much lower than other four tools when applied to all four human ONT datasets (**Table S3**). While DeepMod robustness is comparable to other tools when using 5mC positive control dataset from *E. coli* [33] (**Table S4**). Thus, for clarity, we only keep display the other four tools in **Figure 3**-**5**. Specifically, Nanopolish, Megalodon, and DeepSignal outperformed the other two tools on all datasets (**Figure 3A** and **Table S3**). While Nanopolish, Megalodon, DeepSignal, and Tombo exhibit lower accuracy (less than 0.90) at discordant non-singletons, consistent across four datasets (**Figure 3A, Figure S3A**). Next, we assessed the performance using the ROC curves and AUC in singletons and non-singletons (**Figure 3B**). Again, Nanopolish, Megalodon, and DeepSignal achieved the highest AUC values (singletons AUC: 0.92 - 0.93, non-singletons AUC: 0.96 – 0.98, concordant non-singletons AUC: 0.96 – 0.98, discordant non-singletons AUC: 0.81 – 0.82). We further confirmed the performance assessment using the F1 score (**Figure S3B-S3C**), which is the harmonic mean of precision and recall, and addresses any imbalanced classes. Overall, Nanopolish, Megalodon, and DeepSignal are consistently the top three performers at singleton and non-singleton 5mC prediction at single-read, single-base resolution.

**Figure 3.**
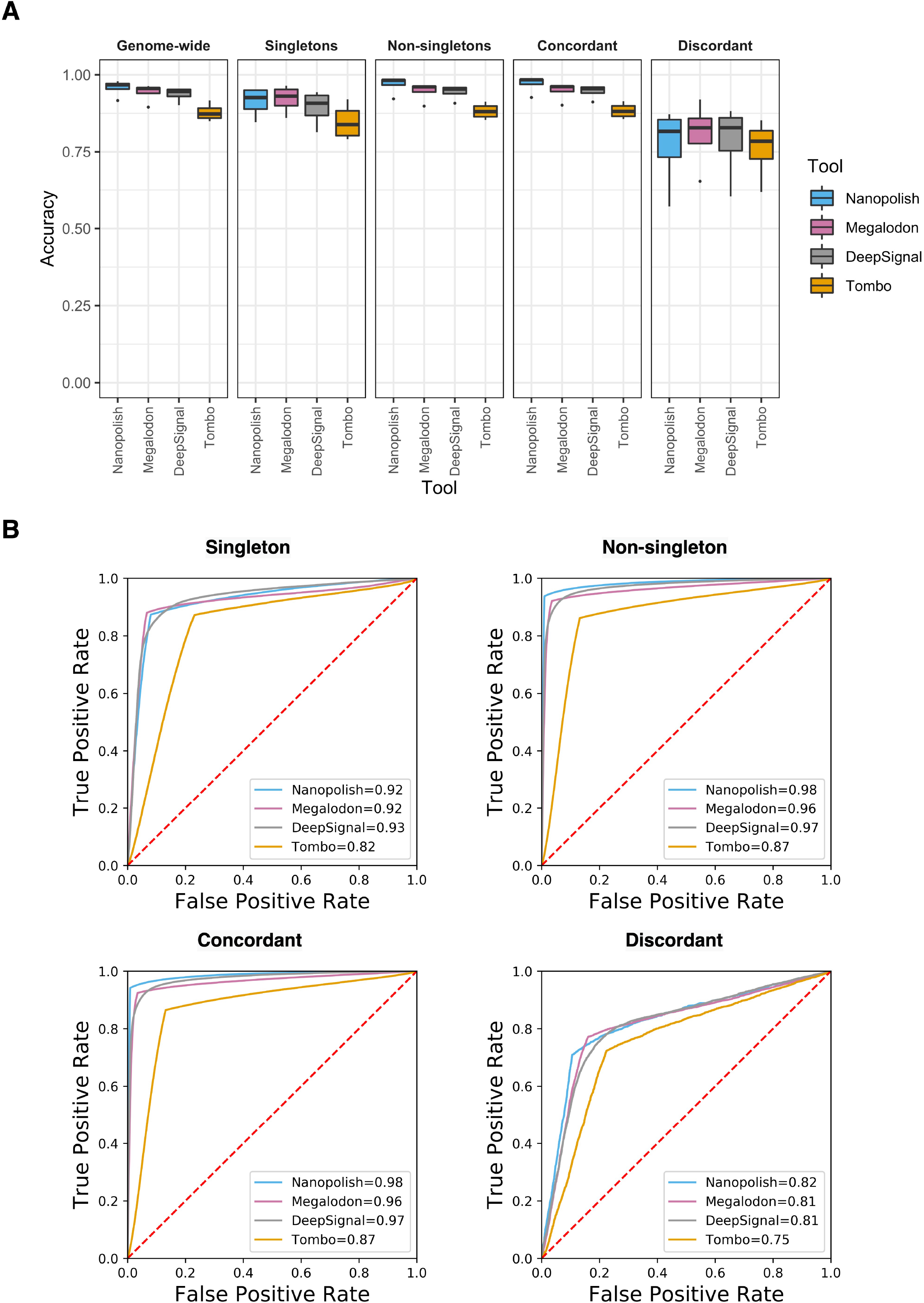
Comparison of Nanopore methylation calling tools for the detection of CpG methylation on four real world data sets in singletons and non-singletons. (**A**) Prediction accuracy across four datasets based on singleton and non-singleton classification. Singletons are CpG sites that contain only one CG within the 10 base pairs, non-singletons are those CpGs that contain more than one CG. Non-singletons in absolute states include concordant non-singletons: all CpGs inside have the same absolute state (i.e., all 100% or all 0% methylated); discordant non-singletons: at least one CpG is fully methylated and at least one other CpG is fully unmethylated. (**B**) ROC curves on NA19240 dataset on singleton, non-singleton, concordant, discordant coordinates.

**Figure 4.**
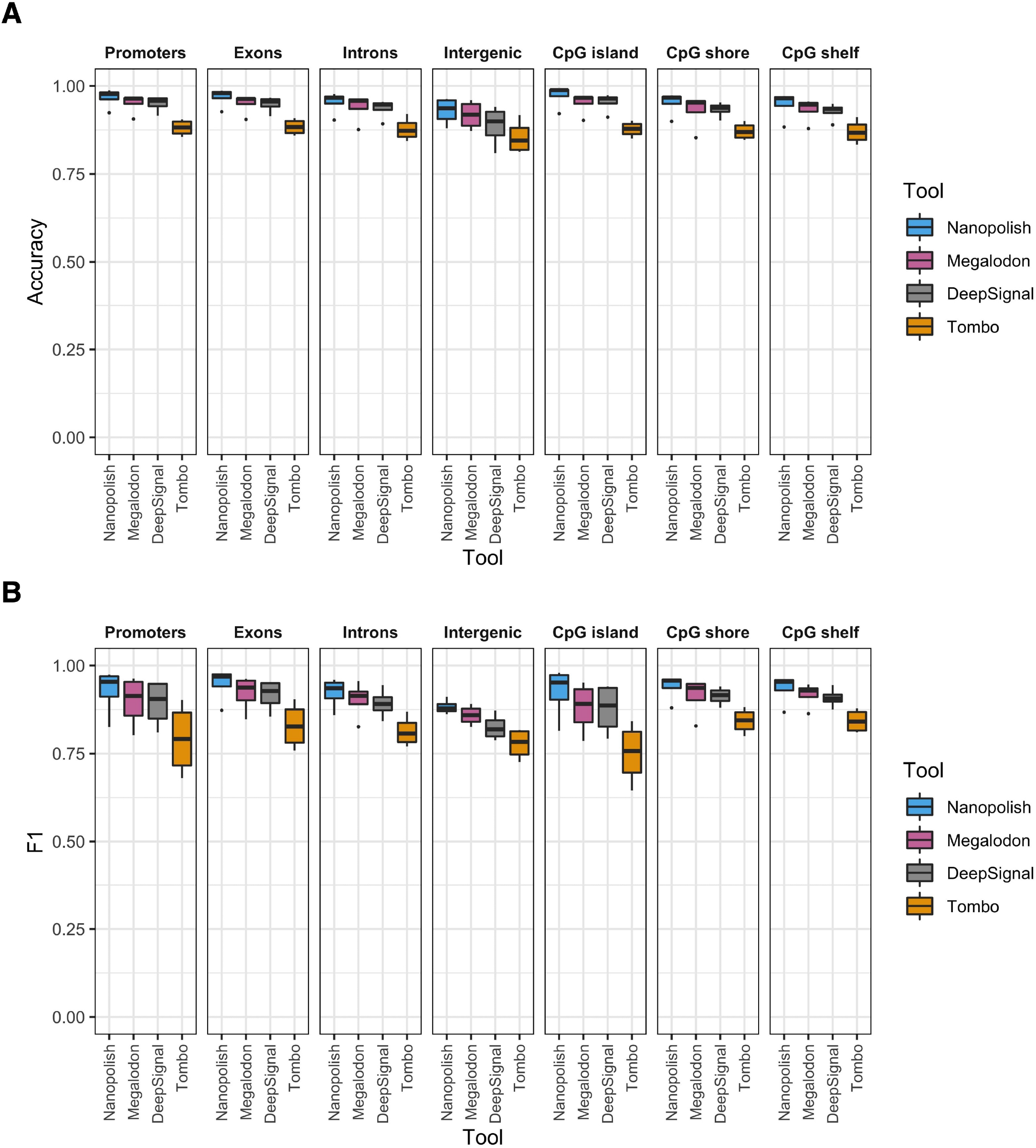
Comparison of Nanopore methylation calling tools for the detection of CpG methylation on four real world data sets in biologically relevant genomic contexts. (**A**) Prediction accuracy across four datasets based on biologically relevant genomic contexts. The biologically relevant genomic contexts include Genome-wide, CpG Islands, promoters, exons, intergenic regions (intergenic) and introns. Promoter is 2000 bp around transcription start site (TSS). (**B**) F1 score across four datasets based on biologically relevant genomic context.

**Figure 5.**
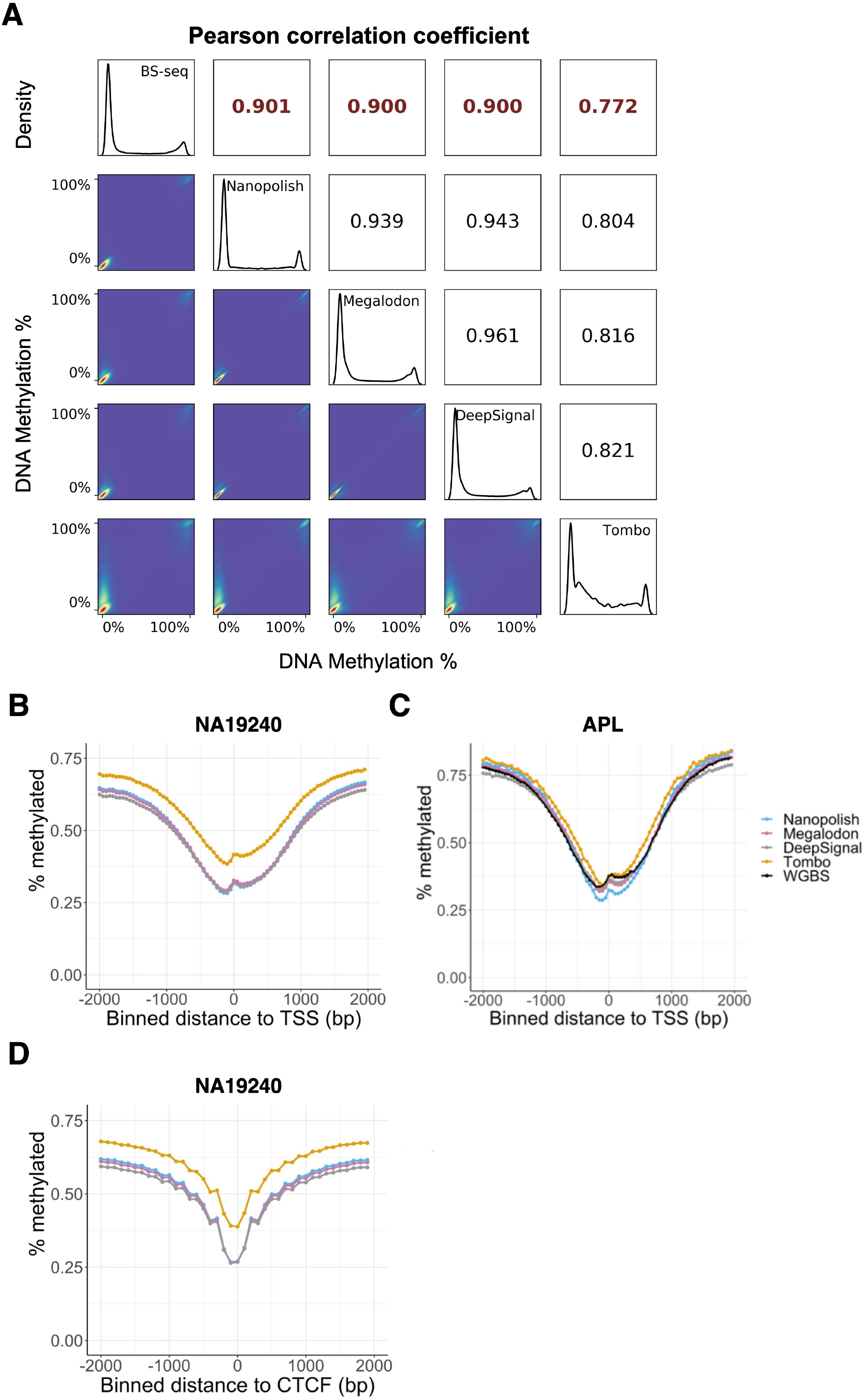
Comparison of Pearson correlation of methylation call tools across all CpG sites. (**A**) Correlation plot showing Pearson correlation of each methylation calling tool with BS-Seq on NA19240. The upper left triangle denotes Pearson’s correlation coefficients; the diagonals are distributions of 5mC percentage of BS-seq data and 5mC percentage predicted by each tool using ONT data. 2D kernel density plots are shown at the lower left triangle for each pair of comparison. (**B-C**) Relationship between CpG methylation percentage and distance to annotated TSS in (**B**) NA19240 and (**C**) APL. (**D**) Relationship between CpG methylation percentage and distance to annotated CTCF binding peaks in NA19240. Distances are binned into (**B, C**) 50-bp, and (**D**) 100-bp windows. Negative distances are upstream and positive distances are downstream of the (**B-C**) TSS and CTCF binding peaks (**D**).

### Predictions of 5mC at single-molecule, single-base resolution across various biologically relevant genomic regions

Since different genomic contexts display various CpG density and DNA methylation levels [43], we overlapped CpG islands, promoters, exons, introns, and intergenic regions (referred as intergenic) with BS-seq (coverage>=5) to evaluate the impact of biologically relevant genomic contexts on 5mC predictions (**Figure 4A-B, Figure S4, Table S3**). Specifically, we define the region 2000 bp around transcription start site (TSS) as the promoter. Nanopolish, Megalodon, and DeepSignal exhibit higher overall accuracy on genome wide CpG sites across all datasets, and overall, intergenic regions display the lowest accuracy (**Figure 4A**). Whereas the overall F1 scores for all tools are not as accurate at CpG island and promoters. Megalodon and DeepSignal are less accurate on CpG islands and promoters (F1 score <0.88) than other regions **(Figure 4B**). The decreased F1 at these two regions may be caused by highly imbalanced distribution of 5mC and 5C on: CpG islands (9,054:166,126) and promoters (11,495:164,989) regions for NA19240 (**Table S3**). In comparison, DeepMod exhibits lower accuracy and F1 score across all genomic regions (**Table S3**). In summary, we concluded that Nanopolish, Megalodon, and DeepSignal achieved better methylation calling performance across genomic contexts.

### Methylation calls by Nanopolish, Megalodon, and DeepSignal show high concordance with ground truth BS-seq

To assess the performance of 5mC prediction of these tools for CpG sites with full range of methylation levels, we evaluated the Pearson’s correlation coefficient between methylation patterns of the predicted DNA methylation percentage (read coverage>=3) and the corresponding BS-seq data (coverage>=5) at single-base resolution. We found that the methylation levels for all CpG sites predicted by Nanopolish, Megalodon and DeepSignal showed highest correlation (**Figure 5A**) with NA19240 reduced representation bisulfite sequencing (RRBS) data. We also observed that the results of Nanopolish, Megalodon, and DeepSignal are highly correlated (R >= 0.94) for NA19240. Similar correlation coefficients of these three tools can be found for APL, K562, and HL-60 (**Figure S5)** datasets. Ideally, as ground truth BS-seq suggested, a bimodal distribution of DNA methylation is expected (0 for unmethylated, 1 for methylated). The histogram of the DNA methylation output of Nanopolish, Megalodon, and DeepSignal displayed bimodal distribution as the BS-seq data. In contrast, Tombo exhibits different data distributions. The DNA methylation level histogram of Tombo output had multiple peaks between 0% and 100% methylation levels. The Pearson’s correlation between BS-seq and DeepMod with (R = −0.07) for NA19240 data indicates that DeepMod cannot effectively predict methylation distribution at whole-genome level for human cells (**Table S5**). We further evaluated the Pearson’s correlation coefficient of methylation percentage achieved by methylation-calling tools with BS-seq across different genomic. Nanopolish, Megalodon and DeepSignal consistently produce the highest correlation coefficients at all genomic regions for NA19240 data (**Table S5 and Figure S6**).

To assess the biological context of the methylation calls, we explored the relationship between CpG methylation percentage and distance to annotated TSS (**Figure 5B-C and S7A-B**). As expected, CpG sites near TSS tend to be unmethylated. Methylation level gets higher as the distance from the TSS increased. DNA methylation patterns from Nanopore sequencing closely resemble the pattern for the WGBS data (**Figure 5C and S7A**). Nanopolish displayed the lowest DNA methylation levels at TSS.

Transcriptional factors CCCTC-binding factor (CTCF) binding sites are featured with low DNA methylation [44]. CTCF plays a critical role in long-range chromatin interactions, the formation and maintenance of the topologically associated domains, and transcription. Thus, we further assessed the relationship between CpG methylation percentage and distance to the center of the CTCF binding peaks from the ChIP-seq data of the matching cell lines (NA19240, K562, and HL-60). Indeed, DNA methylation is the lowest at the center of the CTCF binding peaks (**Figure 5D** and **S7C-D**) and the ONT 5mC calls by Nanopolish, Megalodon, and DeepSignal closely track the pattern of WGBS data (**Figure S7C**).

Overall, Nanopolish, Megalodon, and DeepSignal had high correlations with the background truth BS-seq, and they closely tracked the methylation pattern for the background truth BS-seq at whole genome level. The correlation coefficient of DNA methylation across CpG sites between the five tools and BS-seq is consistent with the read-level accuracy (**Figure 3**-**4**).

### Megalodon and DeepSignal covered more CpG sites than Nanopolish

Lastly, we evaluated the capacity of the five tools to make 5mC prediction for CpG sites by evaluating the number of CpG sites (read coverage>=3) covered by each tool. Megalodon and DeepSignal covered more CpG sites than other tools on four datasets (**Figure 6 and S8, Table S6**). The UpSet diagram shows the number of overlapped sites by the five tools (**Figure 6 and S8**). 52% of the predicted CpG sites in NA19240 were predicted by all five tools (**Table S6**). Furthermore, for all the CpG sites predicted by any of the three top performers (i.e., Nanopolish, Megalodon, and DeepSignal), 92% CpGs were predicted by all three tools, shown by proportional Venn diagram (**Figure 6**). Megalodon and DeepSignal covered more CpG sites that were not covered by the other three tools (Megalodon and DeepSignal predicted 99% of the union of CpG sites using NA19240). Nanopolish covered 93% of the union CpG sites due to the more stringent criteria of log-likelihood ratio used to predict 5mC for non-singletons [9]. Megalodon and DeepSignal covered 6% more CpG sites than Nanopolish and the differences increases greatly for lower sequencing-depth ONT datasets (**Figure 6 and S8, Table S6**). Therefore, Megalodon and DeepSignal predicted the most CpG sites, while Tombo and DeepMod predicted the least CpG sites.

**Figure 6.**
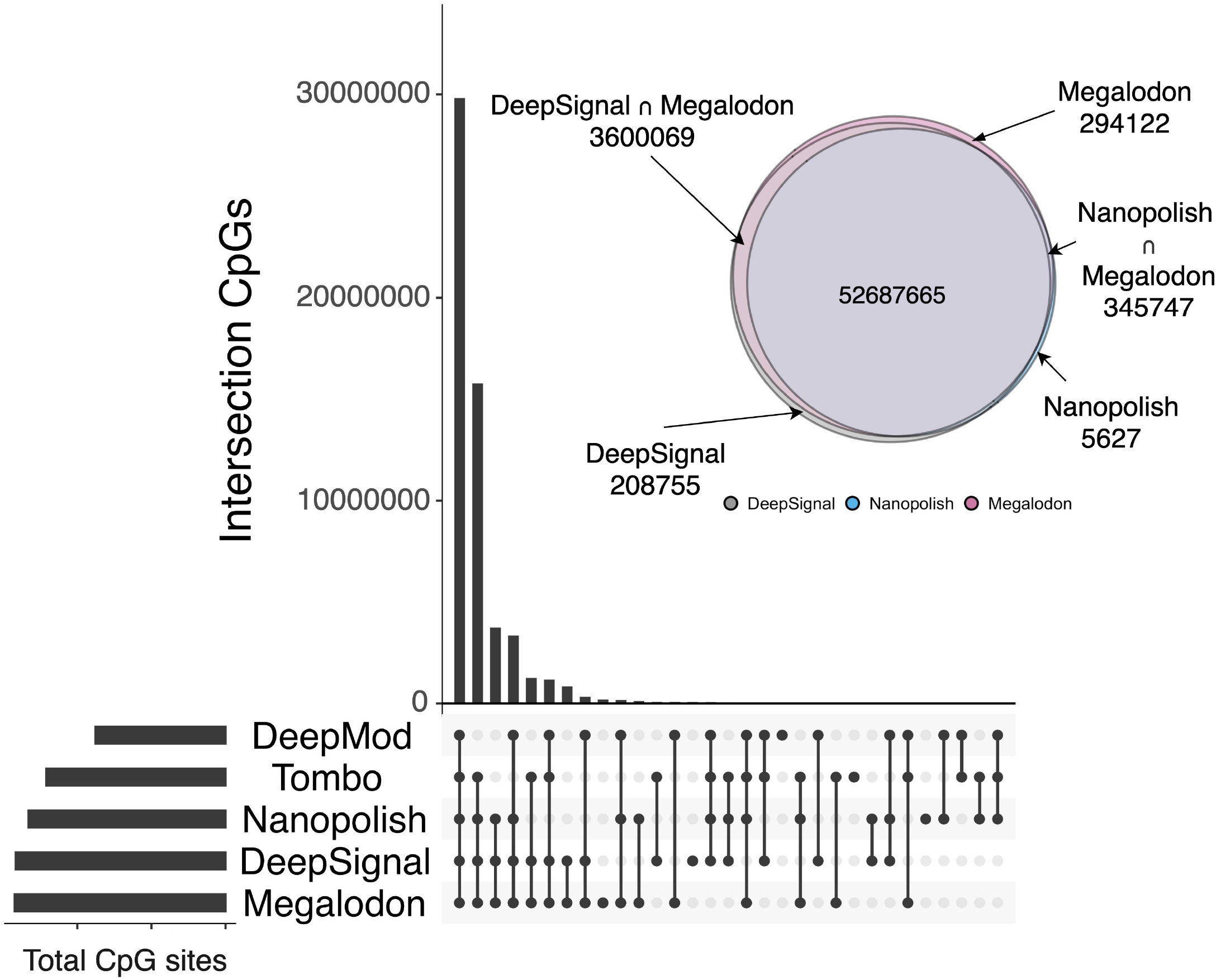
CpG sites detected by methylation calling tools using NA19240. UpSet diagram shown at the lower left is for CpG sites detected by all methylation calling tools. Venn diagram shown at the upper right is for CpG sites detected by top 3 performance methylation calling tools (Nanopolish, Megalodon and DeepSignal). For each methylation calling tool, only CpG sites covered >= 3 reads are considered.

### Running time and memory usage on benchmarking datasets

To evaluate the running time and peak memory of each methylation-calling tool, we ran five pipelines starting from the initial stage of taking input of raw fast5 files to the final output of the read level and genome level prediction results using the same High-Performance Computing (HPC) platform and environment (See **Methods**). In order to parallelize methylation calling, we split the raw reads of the benchmarking dataset, and start 50 running jobs on each part of reads for each methylation-calling tool. A GPU and eight processors of hardware resources were allocated to each job running GPU accelerated computing supported tools (Guppy, Megalodon, DeepSignal and DeepMod) to minimize run time. The SLURM resource and job management system effectively monitor the usage of computing resources on HPC clusters [45]. Therefore, for each tested dataset we ran all jobs managed by SLURM and calculated the sums of run time totals (hours) and the peak memory usage (GB) based on reported logs of SLURM jobs for each pipeline (**Figure 7, Table S7**). Megalodon and Nanopolish had the shortest run times (698 and 703 hours) to process the fast5 raw signal file for NA19240 (32x coverage). While Tombo, DeepSignal, and DeepMod were much longer (9, 32, and 40x longer, respectively) for the same file. Furthermore, Nanopolish required the lowest peak memory usage (∼19 GB) while Megalodon required the highest peak memory usage (22 times). The same analysis of run time and memory usage for other benchmarking datasets also confirmed the ranking for these tools (**Table S7**). In conclusion, Nanopolish requires the least CPU time and the lowest peak memory usage resource. For other tools, there is a trade-off between prediction performance and running resources. Thus, Nanopolish is more appealing for high-coverage mammalian ONT dataset for 5mC prediction.

**Figure 7.**
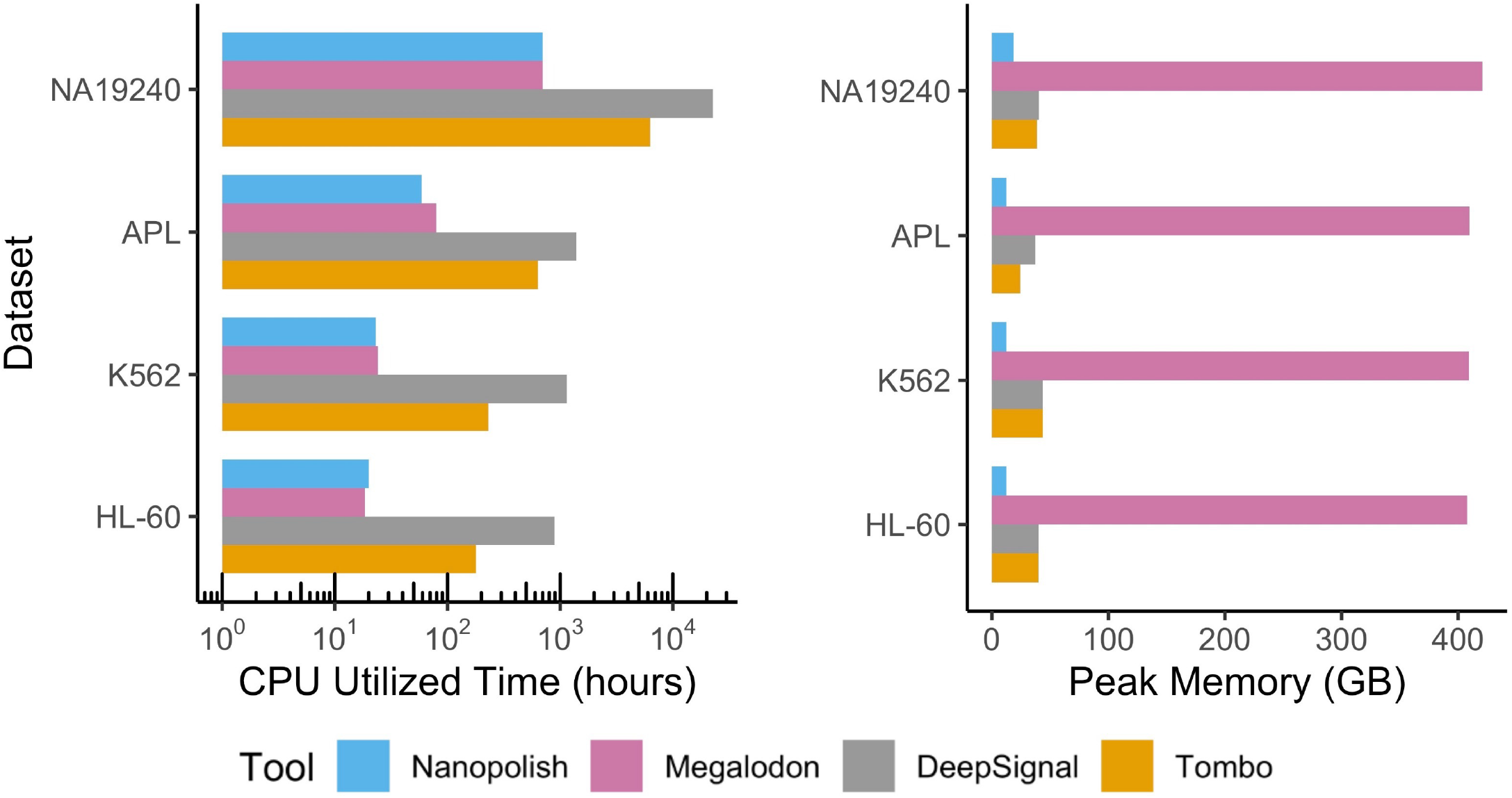
CPU utilized time and memory usage for each methylation calling tool on each dataset. All tools were compared on the same computer clusters: 32 cores, 300 GB RAM, NVIDIA Tesla P100 Data Center and 1 TB Data Direct Networks Gridscalar GS7k GPFS storage appliance. The HPC platform software and hardware specifications are: slurm manager version: 19.05.5, CPU: Intel(R) Xeon(R) Gold 6136 CPU @ 3.00GHz, GPU: Tesla V100-SXM2-32GB.

## Discussion

Enhanced detection of DNA methylation in the human genome is critical to improve our understanding of the functional impacts of epigenetic modifications. Recently, ONT nanopore-based sequencers have made possible direct DNA sequencing to generate long single-molecule reads at base resolution. ONT long-read contributes to the phasing of base modifications with genetic variants, along individual nucleic acids. Therefore, it allows exploration of epigenetic heterogeneity at single-molecule resolution and can improve our ability to detect DNA modification in long range.

ONT has released multiple commercialized platforms and pore-chemistry versions (see timeline in **Figure 1A**). In 2015, ONT released its first commercialized platform, MinION™ [46, 47], a portable device enabling simultaneous sequencing using up to 512 pores, with the capacity to generate up to 30 GB of DNA data [48]. In 2017, ONT introduced a scaled-up platform, GridION™, allowing analysis of up to five MinION flow cells and generation of up to 100GB of data per run [49]. In 2018, ONT introduced the ultra-high-throughput platform PromethION™ with up to 48 flow cells [50], and later offered PromethION24/48 for much larger-scale sequencing [51]. Nanopore sequencing is considered a paradigm among recent sequencing approaches, because of its unique design enabling significant portability and relatively low cost [11, 52].

In the past, the advantages of long-reads and real-time sequencing have made Nanopore sequencing an effective tool to detect genomic and genetics aberrations such as DNA structural variants and RNA alternative splicing events [53]. Nanopore sequencing have demonstrated its powerful capability of detecting structural variation in lung cancer [54, 55], leukemia [56], and neuron disorder [57–59], and it has been applied to clinical samples for molecular etiology or diagnosis of genomic variants relevant disease [57–62]. Meanwhile, Nanopore sequencing of splicing changes has been utilized in cancer research such as breast cancer [63], leukemia [64, 65], and brain tumor [66]. Such research with Nanopore sequencing has improved our understanding of evolutionary process in human diseases.

Nanopore sequencing also provides new opportunity for epigenetic research. For example, Miga et al provides telomere-to-telomere assembly and DNA methylation maps of human X-chromosome [67] using Nanopore sequencing and Ewing et al. developed a new computational tools and long-read nanopore sequencing for transposable element epigenomic profiling [68]. Recently, some efforts have been made to combine Nanopore sequencing and other methods to epigenomics profiling and chromosome structures exploration. For example, Wongsurawat et al utilized Nanopore Cas9-targeted sequencing to simultaneously assess *IDH* mutation status and *MGMT* methylation level in both cell lines and fresh biopsies of diffuse glioma [69]. And, Lee et al developed a new method based on Nanopore sequencing to evaluate CpG methylation and chromatin accessibility simultaneously [70]. Also, several preprint papers utilized Nanopore sequencing to enhance the understanding of epigenetic heterogeneity and mechanism [71–73].

In this study, we benchmarked state-of-art methylation calling tools for Nanopore sequencing data. Based on our systematic comparison analysis, we revealed four key observations. First, the choice of methylation calling critically affects the level of accuracy, F1 score and AUC score on different Nanopore data sets and at different genomic regions. Second, both the HMM model-based Nanopolish and deep learning-based tools Megalodon and DeepSignal, are comparable in terms of overall accuracy, F1 score and AUC values, at single-base and single-read resolution. Notably, Nanopolish has the lowest memory usage, and both Nanopolish and Megalodon are faster than DeepSignal. Third, the methylation detections in discordant regions with mixed DNA methylation and intergenic regions exhibit lower accuracy and F1 score across all five tools. Nanopolish is fast and accurate, at the same time, it outputs the methylation levels of 6% fewer CpG sites than DeepSignal and Megalodon, due to the more stringent log-likelihood ratio cutoff for predicting non-singleton CpG sites. Nanopolish can be used for quick prediction, and future algorithm development can focus on increasing the accuracy and higher CpG coverage, which leads to higher overall performance. When high-performance clustering or cloud computing is available, Nanopolish, Megalodon, and DeepSignal can each produce high-quality methylation predictions on the largest number of CpG sites. In the absence of an HPC or cloud it is feasible to run Nanopolish on a laptop for DNA methylation calling due to its short run-time and low memory for in-field analysis that also makes it compatible with ONT MinION’s portability.

We believe that our benchmarking of methylation calling tools will guide researchers and practitioners to make conscious and effective choices when designing the analytic plan for epigenomic profiling using ONT sequencing, including Nanopore Cas9-targeted sequencing data analysis. The bottlenecks revealed by our analysis can help developers to improve ONT sequencing data methylation-calling training data generation and tool design. We note that one recent preprint [33] proposed a consensus random forest model to improve accuracy by combining read level methylation predictions of some tools (i.e., DeepSignal and Megalodon). Our analysis demonstrates that a training dataset covering discordant non-singletons and intergenic regions would improve the overall robustness of DNA methylation prediction at single-molecule, single-base resolution for human epigenome-wide study.

## Conclusion

Oxford Nanopore long-read sequencing technology poses a challenge for accurate methylation predictions. The past few years have witnessed rapid development of both the sequencing technology and analytical tools. For DNA methylation analysis, many algorithms are emerging for ONT sequencing data. We comprehensively surveyed current publicly available computational tools for direct ONT DNA sequencing data methylation detections. We systematically evaluated the advantages, disadvantages, and identified performance bottlenecks that affect the robustness of DNA methylation detection at single-molecule and single base resolution. Using a standardized workflow we assessed the performance of five DNA methylation calling tools and found that methylation callers vary in their accuracy in diverse genomic contexts, epigenome coverage, peak memory usage, and run time, for both single-read and single-base resolution. For initial DNA methylation analysis, we recommend Nanopolish given its short run-time, low memory requirement and overall high performance in calling DNA methylation in whole genome level, singleton, non-singleton, promoter, CpG islands, exonic, and intronic regions. For systematic analysis, we recommend integrating the preliminary results with Megalodon, or DeepSignal output for optimal performance, e.g., more comprehensive epigenome coverage. Comprehensive and balanced training datasets that cover various genomic contexts is desirable for more robust prediction of DNA methylation in discordant and intergenic regions and will help improve our understanding of epigenetic mechanisms underlying many different biological processes, such as aging and cancer development.

## Methods

### Sample collection and processing

In the study we provided four independent human datasets - one normal B-Lymphocyte cell line (NA19240) [39], one primary acute promyelocytic leukemia clinical specimen (APL), two cancer cell lines (K562, HL-60).

For APL, sample was obtained from the Stem Cell and Xenograft Core of the University of Pennsylvania. The Core maintains a tissue bank of cells from patients with Hematologic Malignancies. This is Institutional Review Board (IRB) approved research (IRB protocol #703185). The patient sample was collected at the time of clinical presentation and prior to therapy. The sample was collected as leukopheresis and viably frozen using standard techniques. The de-identified specimen was then provided to the Jackson Laboratory for Genomic Medicine (JAX-GM). Diagnosis (Dx) of acute promyelocytic leukaemia (APML) was confirmed by Fluorescence in situ hybridization (FISH) analysis for t(15;17). K562 and HL-60 were cultivated in Roswell Park Memorial Institute (RPMI) 1640 Medium (Gibco, A10491-01) with 10% fetal bovine serum (FBS) (Gibco, 26140079). K562 medium was additionally supplemented with 1% Antibiotic-Antimycotic (Gibco, 15240062). HL-60 medium was additionally supplemented with 1.2% of penicillin-streptomycin (Gibco, 15140-163), GlutaMAX (Gibco, 35050-061), Sodium Pyruvate (Gibco, 11360-070), MEM Nonessential Amino Acids (Corning, MT25025CI) and MEM Vitamin Solution (Corning, MT25020CI). Incubator conditions were 37°C and 5% CO_2_.

### Bisulfite sequencing (BS-seq) dataset and analysis

We generated whole genome bisulfite sequencing (WGBS) for APL. DNA was extracted using AllPrep DNA/ RNA kit (Qiagen) following manufacturer’s recommendation. Two 500ng DNA were sheared to 500bp using a LE220 focused-ultrasonicator (Covaris) and purified using 0.9X SPRI beads (Beckman Coulter). The libraries were prepared using the KAPA Hyper Prep Kit for Illumina (Roche) and bisulfite conversion was performed using the TrueMethyl Seq Kit (CEGX). Briefly, the fragmented DNA was first spiked in with CEGX sequencing controls, followed by end-repair and A-tailing, and then ligated with SeqCap indexed adaptor (Roche). Sample destined for 5hmC library was first subjected to oxidation whereas samples destined for 5mC library was treated as mock. This is then followed by a bisulfite conversion. The treated DNA were cleaned up and amplified with 15 cycles of PCR and purified. The final library was quantified by real time qPCR for an accurate concentration. Libraries were sequenced paired end 2×150bp on the Illumina HiSeq 2500 instrument.

We utilized the published whole genome bisulfite sequencing (WGBS) for K562 (ENCODE accession number: ENCFF721JMB, ENCFF867JRG), and reduced representation bisulfite sequencing (RRBS) for HL-60 (ENCODE accession number: ENCFF000MDA, ENCFF000MDF) and NA19240 (ENCODE accession number: ENCFF000LZS, ENCFF000LZT).

All BS-seq data were analyzed with Bismark [74] with the human reference genome (GRCh38/hg38) to get the cytosine methylation frequency at each CpG site. Region-specific analysis and local smoothing for samples was performed using the BS-seq package (https://github.com/TheJacksonLaboratory/BS-seq-pipleine). Then, we only select high-confidence CpG sites with coverage >=5, where a CpG is considered as fully methylated when its methylation frequency is 100% and considered as unmethylated when its methylation frequency is zero (**Table S2**). In total, 30,377 singleton CpGs (10,815 fully methylated and 19,562 unmethylated) and 224,645 non-singleton CpGs (29,432 fully-methylated and 195,213 unmethylated) were selected from NA19240 RRBS, and 42,137 singleton CpGs (21,738 fully-methylated and 20,399 unmethylated) and 276,411 non-singleton CpGs (56,444 fully-methylated and 219,967 unmethylated) were selected from HL-60 RRBS. For K562 and APL, the total selected high-confidence CpG sites are 25,382,453 and 8,707,630 respectively from WGBS. For each dataset, we take the intersections of high-confidence sites from all tools and BS-seq as our final high-confidence set and the selected high-confidence sites **(Table S3)**.

### Nanopore sequencing dataset and analysis

We generated Nanopore sequencing dataset for APL, K562, and HL-60 at JAX-GM. For APL, genomic libraries were prepared using the Rapid Sequencing Kit (SQK-RAD004, ONT) according to manufacturer’s recommendation. Briefly 1200ng DNA was incubated with 2.5ul of FRA at 30°C for 1 min and 80°C 1 min. This is followed by an addition of 3ul of adaptor (RAP) to the reaction mix and incubated at 5 min at room temperature. The libraries were sequenced on the flowcell R9.4.1 (FLO-MIN106, ONT) on GridION (ONT) using the MinKNOW software for 48hr.

For K562 and HL-60, HMW genomic DNA were extracted from 5m cells using phenol chloroform approach (PMID30933081). Libraries were prepared using the Rapid Sequencing Kit (SQK-RAD004, ONT) according to manufacturer’s recommendation. Briefly 1200ng DNA was incubated with 2.5ul of FRA at 30°C for 1 min and 80°C 1 min. This is followed by an addition of 3ul of adaptor (RAP) to the reaction mix and incubated at 5 min at room temperature. The libraries were sequenced on the flowcell R9.4.1 (FLO-MIN106, ONT) on a GridION (ONT) using the MinKNOW software for 48hr.

For NA19240, we request Nanopore raw data from previously published research [39]. For *E.coli*, we utilized an example dataset on Github (https://github.com/comprna/METEORE/tree/master/data/example), which contains 50 single-read fast5 files from the positive control dataset for *E.coli* generated by Simpson et al [9].

All Nanopore reads (.FAST5 files) were base-called by Guppy (v4.2.2) with default high-accuracy model (dna_r9.4.1_450bps_hac.cfg). The base-called reads were then aligned to human reference genome (GRCh38/hg38) for human dataset (NA19240, APL, K562, HL-60) or aligned to the *E.coli* K12 MG1655 genome for *E.coli* dataset using minimap2 [41]. Specially, R9.4-series pore is the current broadly used Nanopore flow cell and there is a slight difference between R9.4 and R9.4.1 flow cells and most computational model can work for both [75].

Additionally, for cell line authentication of K562 and HL-60, we aligned the base-called reads to the human reference genome (GRCh37/hg37) with the help of Minimap2 [41] and Samtools [76] and compared target regions in aligned reads with reported insertions/deletions (indels) derived from the Cancer Cell Line Encyclopedia (CCLE) project [77] in genome browser IGV to identify the cancer cell line information.

### Experimental settings and running configurations for Nanopore sequencing analysis

We ran five tools on the benchmarking datasets. We supply FAST5 files generated by Nanopore sequencers with raw signals and base calls as input for methylation detection analysis. To compare speed performance, all tools were carried out on the same computer clusters: 32 cores, 300 GB RAM, NVIDIA Tesla P100 Data Center and 1 TB Data Direct Networks Gridscalar GS7k GPFS storage appliance. The HPC platform software and hardware specifications are: slurm manager version: 19.05.5, CPU: Intel(R) Xeon(R) Gold 6136 CPU @ 3.00GHz, GPU: Tesla V100-SXM2-32GB.

Base calling, the process of translating raw electrical signal of the sequencer into nucleotide sequence, is the initial step of Nanopore data analysis. Both ONT and independent researchers are actively developing different tools for base calling step. Specifically, ONT provides base-calling programs including official ONT community-only software (Albacore and Guppy) and open-source software (Flappie, Scrappie, Taiyaki, Runnie, and Bonito), the latter of which are under development with new algorithms for base calling [31, 78, 79]. Only very recently has it been possible to base-call DNA modifications directly from the raw signal without genomic anchoring, which can be accomplished via specific base callers such as Scrappie [80]. Among the base calling programs, Albacore and Guppy are compatible with Nanopore R9.4 reads and offer the most stable performance [42]. Albacore [81, 82] is a general-purpose base caller that runs on CPUs. Guppy [83] is a neural network based basecaller with several bioinformatic post-processing features. Guppy supports both CPUs and GPUs for improved base-calling run time, and it is available on the ONT community site (https://community.nanoporetech.com) for internal use. Because the state-of-art basecaller Guppy using the default model showed excellent performance among ONT basecalling tools [42], we utilized Guppy (v4.2.2, with all 32 CPU threads) for base calling for all datasets and all DNA methylation calling tools.

RNN and HMM are computationally intensive algorithms. In HMM-based Nanopolish tool, the Viterbi algorithm is used for methylation prediction. The Viterbi algorithm is a sequential technique, and its computation cannot currently be parallelized with multithreading. However, in RNN-based DeepSignal and DeepMod, multiple threads can work on different sections of the neural network and thus RNN computation can be parallelized with multithreading. We choose this system for evaluation since it has a larger memory capacity than desktop systems and, with the help of a large number of cores, the tasks can be easily parallelized to accelerate data output for state-of-the-art tools.

### Methylation calling Nanopore sequencing at read level and site level

We evaluated the performance of Nanopolish (v0.13.2), Megalodon (v2.2.9), DeepSignal (v0.1.8), Tombo (v1.5.1), and DeepMod (v0.1.3) to detect 5mC at CpG dinucleotides. These five tools differ in the underlying algorithms and the modifications they are trained to detect.

Nanopolish [9] calls 5mC in a CpG context using a HMM to assign a log-likelihood ratio (LLR) for each CpG site, where a positive log-likelihood ratio (LLR) indicates support for methylation. Nanopolish groups nearby CpG sites together and calls the cluster jointly to assign the same methylation status to each site in the group. For example, on a motif such as CGCGT, Nanopolish reports a LLR for the whole group, rather than a separate LLR for the individual cytosine. We use the 2.0 as the LLR threshold for methylation calling as the Nanopolish authors suggests that the initial 2.5 shown in the paper is overly conservative, and the threshold was replaced with 2.0 from v0.12.0 [84]. To be more specific, we first called methylation at the read level: we removed ambiguous reads when the absolute value of their LLR was less than 2.0, and then called CpG sites as methylated when the LLR > 2.0 and called CpG sites as unmethylated when the LLR < −2.0. Then we calculated methylation frequency at the site level by converting the LLR to a binary call (methylated/unmethylated) for each read and calculating the fraction of reads classified as methylated.

Megalodon [24] is a new ONT-developed a research command line tool and can identify modified base and sequence variant calls from raw nanopore reads. For modified base calls, Megalodon utilizes Guppy (v>=4.0) on the backend and pre-trained models for basecalling. It anchors the intermediate basecalling neural network output to a reference genome. Megalodon performs the methylation calling at either the per-read or per-site level (aggregate per-read results) based on the log probability that the base is modified or canonical. Guppy (v>=4.0) backend and pre-trained models is recommended for base calling, so we fed Megalodon with Guppy v4.2.2 with the latest 5mC in an all context model (res_dna_r941_min_modbases_5mC_v001.cfg) from Rerio [85] as the basecalling model, and chose the default 0.8 threshold as the probability cutoff to count the called base (modified or canonical) with probability >0.8 toward the final aggregated output at per-site level.

DeepSignal [27] proposed a deep recurrent neural network with Bidirectional Long short-term memory (BiLSTM)+Inception structure to detect the methylation state of target cytosine in CpG context. DeepSignal required an extra the re-squiggle module of Tombo before methylation calling. The methylation calling output of DeepSignal is a tab-delimited text file (tsv) at read level including two probability values for each base, one for methylated (prob_1) and one for unmethylated (prob_0), as well as a binary call (unmethylated/methylated) for each base. The CpG sites is called as methylated when prob_1 > prob_0 and is called as unmethylated when prob_1 <= prob_0. We performed per-read methylation calling with the CpG model trained using HX1 R9.4 1D reads (model.CpG.R9.4_1D.human_hx1.bn17.sn360.v0.1.7+.tar.gz) provided with the latest version of DeepSignal, and calculated the fraction of reads classified as methylated at site level with their official methylation frequency script.

ONT-developed Tombo [21] performed a statistical test to identify modified nucleotides with its alternative model without the need for prior training data. Tombo computed per-read, per-genome location test statistics by comparing the signal intensity difference between modified bases and canonical bases. We chose to use the recommended CpG motif specific model with the default threshold of (−1.5, 2.5) for DNA where scores below −1.5 were considered as methylated and above 2.5 unmethylated, and scores between these thresholds did not contribute to the per-site methylation. After that, we calculated methylation percentage at each genomic position.

DeepMod [25] designed a bidirectional recurrent neural network (RNN) with an LSTM unit for genome-scale detection of DNA modifications. The input is a reference genome and FAST5 files with raw signals and base calls, and the output is a BED file with coverage, number of methylated reads, and methylation percentage information for genomic positions of interest. Since 5mC in CpG motifs has a cluster effect in the human genome [25], DeepMod provides a cluster model to generate a final output for site level predicted methylation probability in human genome. We performed DeepMod for methylation calling with the RNN model (rnn_conmodC_P100wd21_f7ne1u0_4) and cluster model (na12878_cluster_train_mod-keep_prob0.7-nb25-chr1) [86]. Also, since DeepMod aggregated methylation callings results into a per-site output BED file, we counted the number of methylated callings and unmethylated callings from BED outputs to evaluate its read level performance.

The performances of these methods that use prior knowledge about the expected deviations in signal depend notably on the training data used, which is typically composed of a fully unmodified and a fully modified sample. Motifs that are not represented in the training set or that contain mixtures of modified and unmodified bases lead to suboptimal performance.

### Methylation calling performance evaluation at read level

We designed the performance evaluation process for 5-methylcytosine status prediction among five methylation calling tools.

First, we evaluated performance for the five tools on four real world Nanopore Sequencing datasets at singleton and non-singleton site levels and biologically relevant genomic context level at read level. To be more specific, we only considered CpG sites covered by >= 5 reads in BS-seq and CpGs sites covered by >= 1 reads by methylation calling tools, and joined the common sites identified by the five tools with background truth BS-seq. For the CpG sites that showed 0% or 100% methylation level, we evaluated the performance of these tools as a per-read classification model. Then we joined each tool’s prediction results to a common CpG set and measured accuracy on basis of singleton and non-singleton sites, or biologically relevant genomic context. We compared the percentage of methylation calculated by the five Nanopore-based methods to that derived by BS-seq at annotated locations. On each location basis, we calculated the F1 score, accuracy, precision, recall, and assessed the tradeoff between true-positive and false-positive rates of 5mC prediction by calculating receiver operating characteristic (ROC) curve by varying the threshold for methylation calling and reported the area under the ROC curve (AUC) values. Metrics of performance are calculated as following using BS-seq as ground truth:

**TP**: true positive
**TN**: true negative
**FP**: false positive
**FN**: false negative
**Precision**: TP/(TP+FP)
**Recall**: TP/(TP+FN)
**Accuracy**: (TP + TN) / (TP + TN + FP + FN)
**F1 Score**: 2*(Recall * Precision) / (Recall + Precision). We calculated F1 score for both 5mC and 5C and used macro_F1(average F1_5mC and F1_5C) for final F1 score.
**ROC AUC:** the area under receiver operating characteristic curve, usually ranging from 0.5 to 1.0. It is a performance metric used to evaluate how a classifier performs on both methylated and unmethylated class predictions.

### Methylation calling performance evaluation at site level

We calculated the Pearson’s coefficient between predicted methylation status and BS-seq status and checked the methylation distribution structure for each tool at a genome level. Again, we first kept CpGs with >= 5 reads in BS-seq and CpGs with >= 3 reads by methylation calling tools, joined the CpG sites as overlapped sets, and calculated methylation frequencies for all DNA CpGs at a genome level for each tool from read level. For correlation analysis, we treat each pair of tools as a regression model to calculate Pearson’s correlation coefficients at a genome level. We compute the relationship between CpG methylation percentage with distance to annotated transcription start site (TSS) and transcriptional factors CCCTC-binding factor (CTCF) binding sites using deepTools [87].

### Memory usage and running time for methylation tools

We compared the capability of the five methylation calling tools for memory usage and running time on single-read fast5 file in each dataset. All tools have support for multi-processors, so we generated simulated data sets to compare the scalability of these tools on the same system configurations: We split each dataset into 50 batches to ensure the same scale of input, chose default tool configurations to run the program. Each job was allocated eight processors (200GB per processor) and one GPU hardware resource (GPU is allocated for running Guppy, Megalodon, DeepSignal and DeepMod). We extract running time (field name: CPU Utilized) and peak memory utilization (field name: Memory Utilized) from the SLURM job log data. These results were used as the measurement of running time and memory usage for hardware performance comparison and evaluation.

### CpG sites comparison by different methylation calling tools in each dataset

We compared the number of CpG sites covered by different methylation calling tools. For each dataset, we kept CpGs with >= 3 reads by methylation calling tools, then joined the CpG sites with BS-seq to check the overlapping sites that were also detected in BS-seq. We also checked the CpG sites detected by Nanopolish, Megalodon and DeepSignal since these three performed best among the methylation calling tools.

### Availability of data and materials

All source codes are publicly available at GitHub https://github.com/liuyangzzu/nanome and https://zenodo.org/record/4730517 with the DOI: 10.5281/zenodo.4730517. APL WGBS and ONT datasets, HL-60 and K562 ONT datasets were deposited under GEO accession GSE173675, GSE173676, GSE173687, and GSE173688.

## Acknowledgements

S.L. is supported by a startup fund from The Jackson Laboratory for Genomic Medicine, Leukemia Research Foundation New Investigator Grant, The Jackson Laboratory Director’s Innovation fund 19000-17-31 and 19000-20-05, The Jackson Laboratory Cancer Center New Investigator Award, The Jackson Laboratory Cancer Center Fast Forward Award, and the National Institute of General Medical Sciences of the National Institutes of Health under Award Number R35GM133562. Research reported in this publication was partially supported by the National Cancer Institute of the National Institutes of Health under Award Number P30CA034196. The content is solely the responsibility of the authors and does not necessarily represent the official views of the National Institutes of Health. The authors thank Dr. Charles Lee for scientific discussion and providing the access to the long-read sequencing data. The authors thank Dr. Derya Unutmaz for helpful discussion. The authors thank the members of Li Lab for discussions and thank Drs. Stephen Sampson and Kevin Seburn from The Jackson Laboratory Research Program Development for editing this paper. The authors thank Dr. Chia-Lin Wei and Dr. Chew Yee Ngan from Genome Technologies at and The Jackson Laboratory for Nanopore sequencing support and discussion. The authors thank The Jackson Laboratory Computational Sciences and Research IT team for technical support, e.g., Shiny app deployment helped by Sandeep Namburi.

## Declarations

None of the authors have any competing interests.

## SUPPLEMENTARY INFORMATION

**Additional file 1:** Supplementary figures. This file contains supplementary figures S1-S8.

**Additional file 2:** Supplementary tables. This file contains supplementary tables S1-S7.

