## Supplementary Figures for "DNA methylation calling tools for Oxford Nanopore sequencing: a survey and human epigenome-wide evaluation"

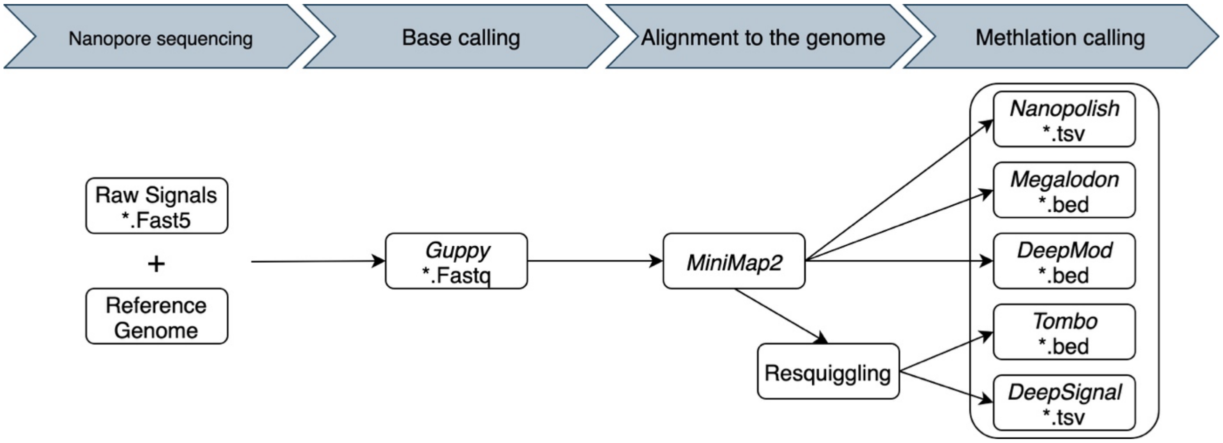


**Figure S1: Workflow for 5-methylcytosine (5mC) detection for Nanopore sequencing.** The pipeline has four steps: (1) Nanopore sequencing. (2) Base calling, which requires raw signals and reference genome as input to perform base calling by Albacore or Guppy. (3) Alignment to the genome by direct mapping with miniMap2 and re-squiggle with Tombo (optional). (4) Methylation calling. Here we compare five methylation calling tools, Nanopolish, Megalodon, DeepSignal, Tombo, and DeepMod to detect cytosine status in CpG context.

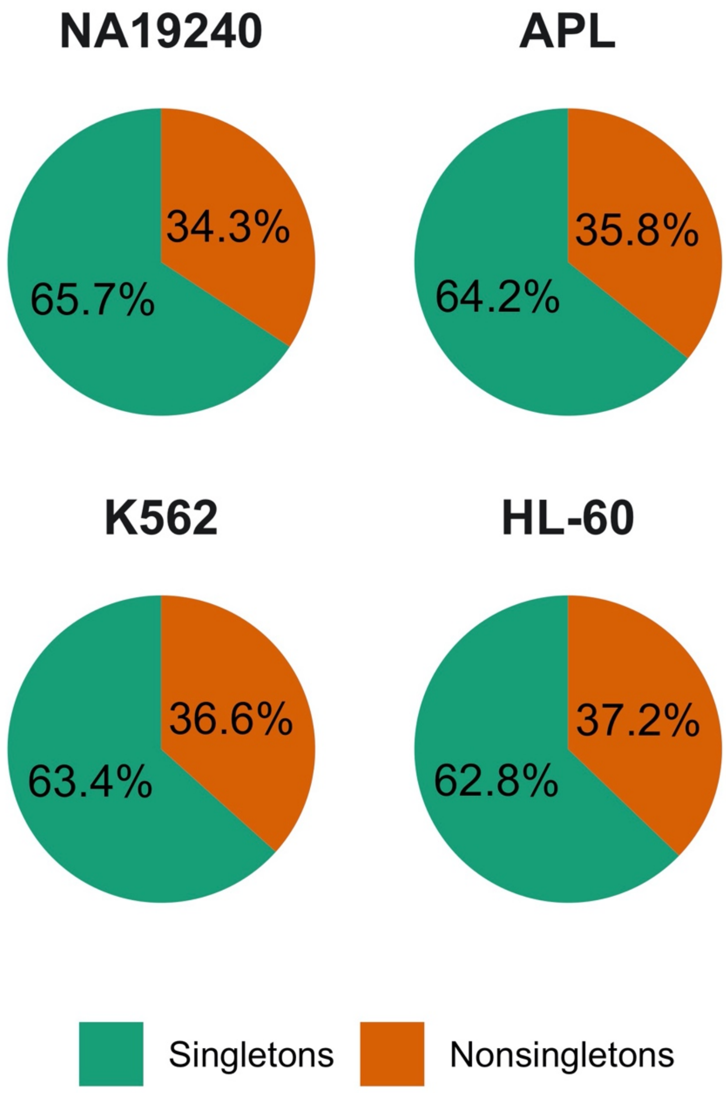


**Figure S2: Distribution of CpG sites at singleton and non-singleton regions covered by raw Nanopore reads (read coverage>=3).**

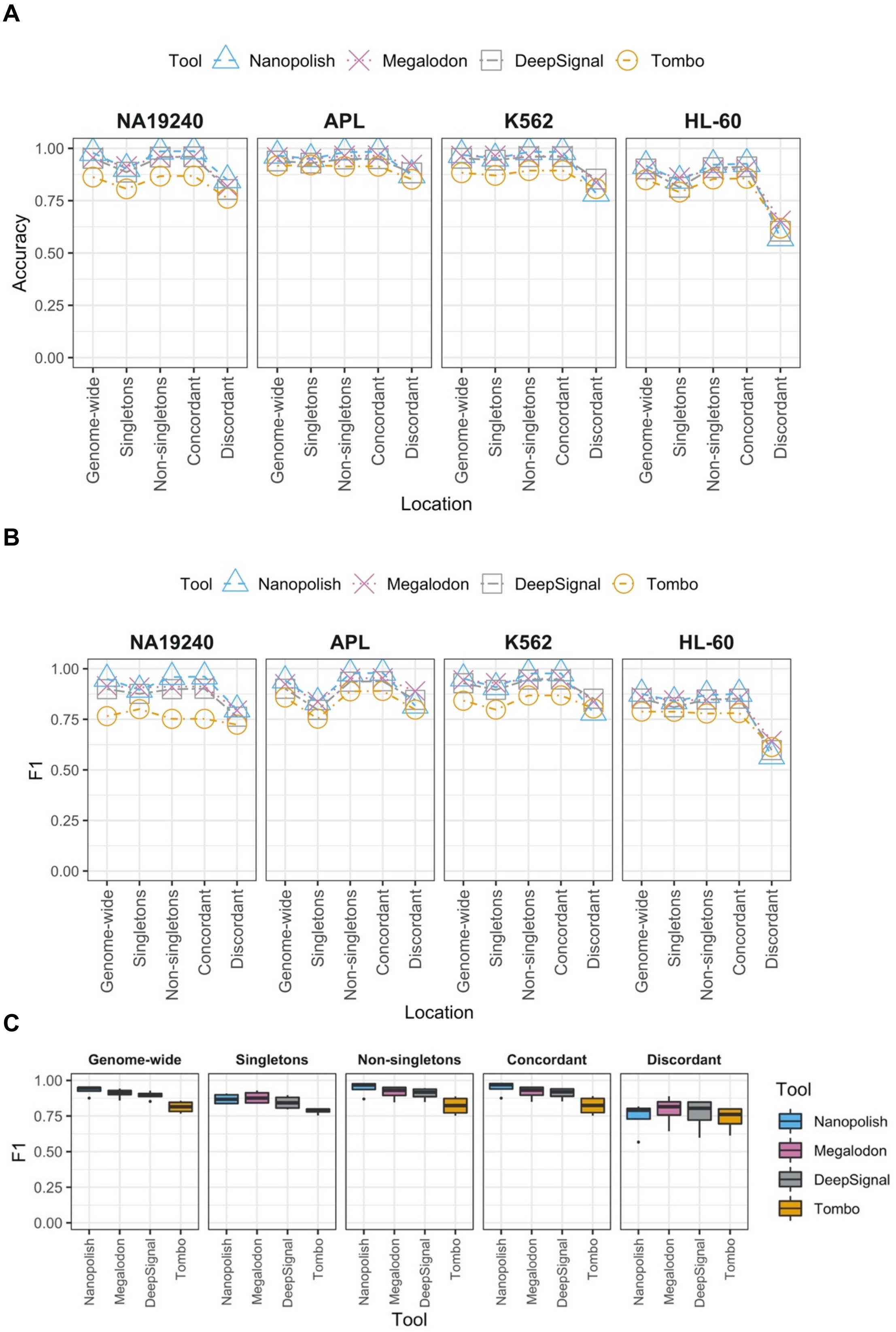


**Figure S3: Accuracy (A) and F1 score (B-C) on four datasets across singletons and non-singletons.**

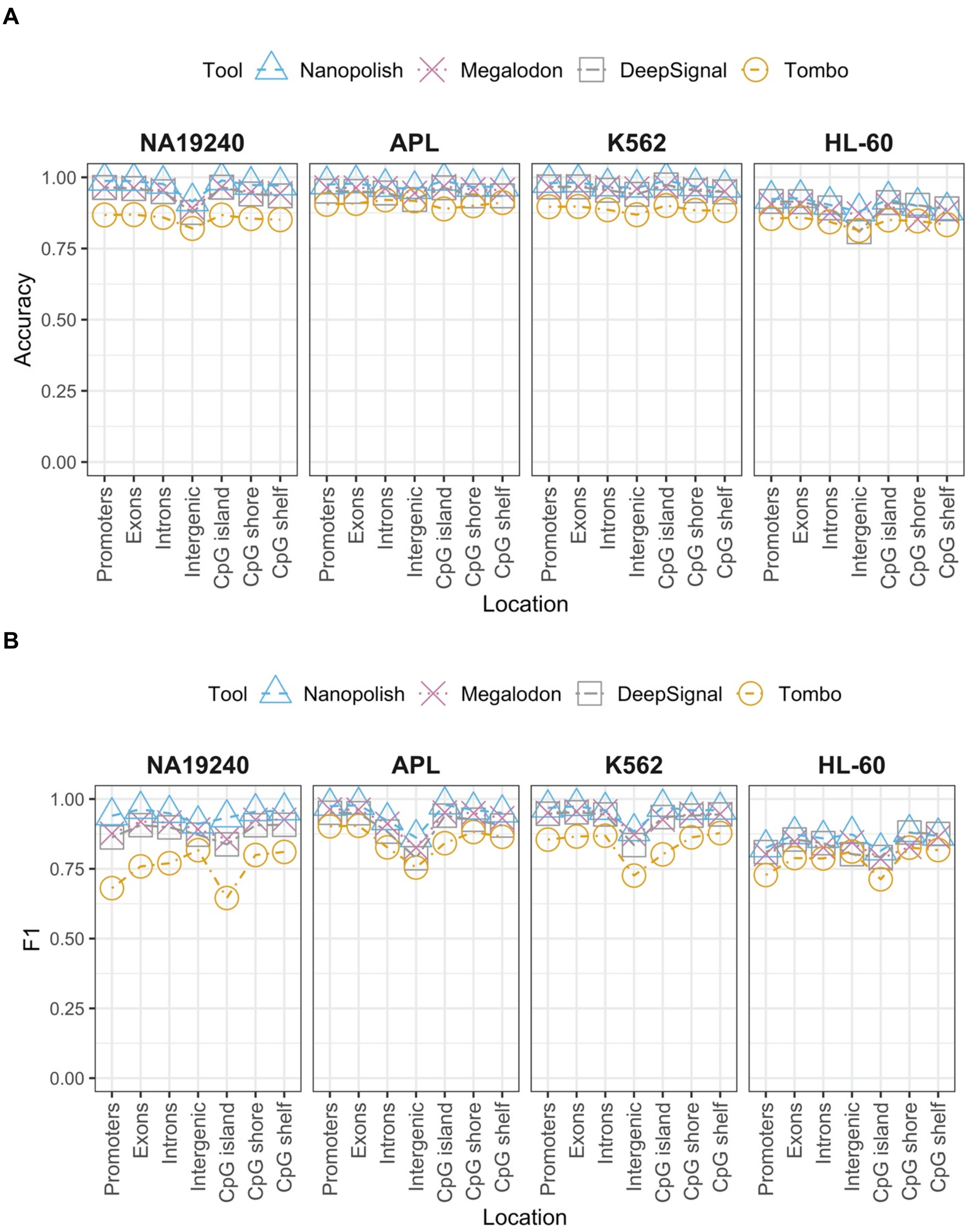


**Figure S4: Accuracy (A) and F1 score (B) on four datasets across different genomic regions.**

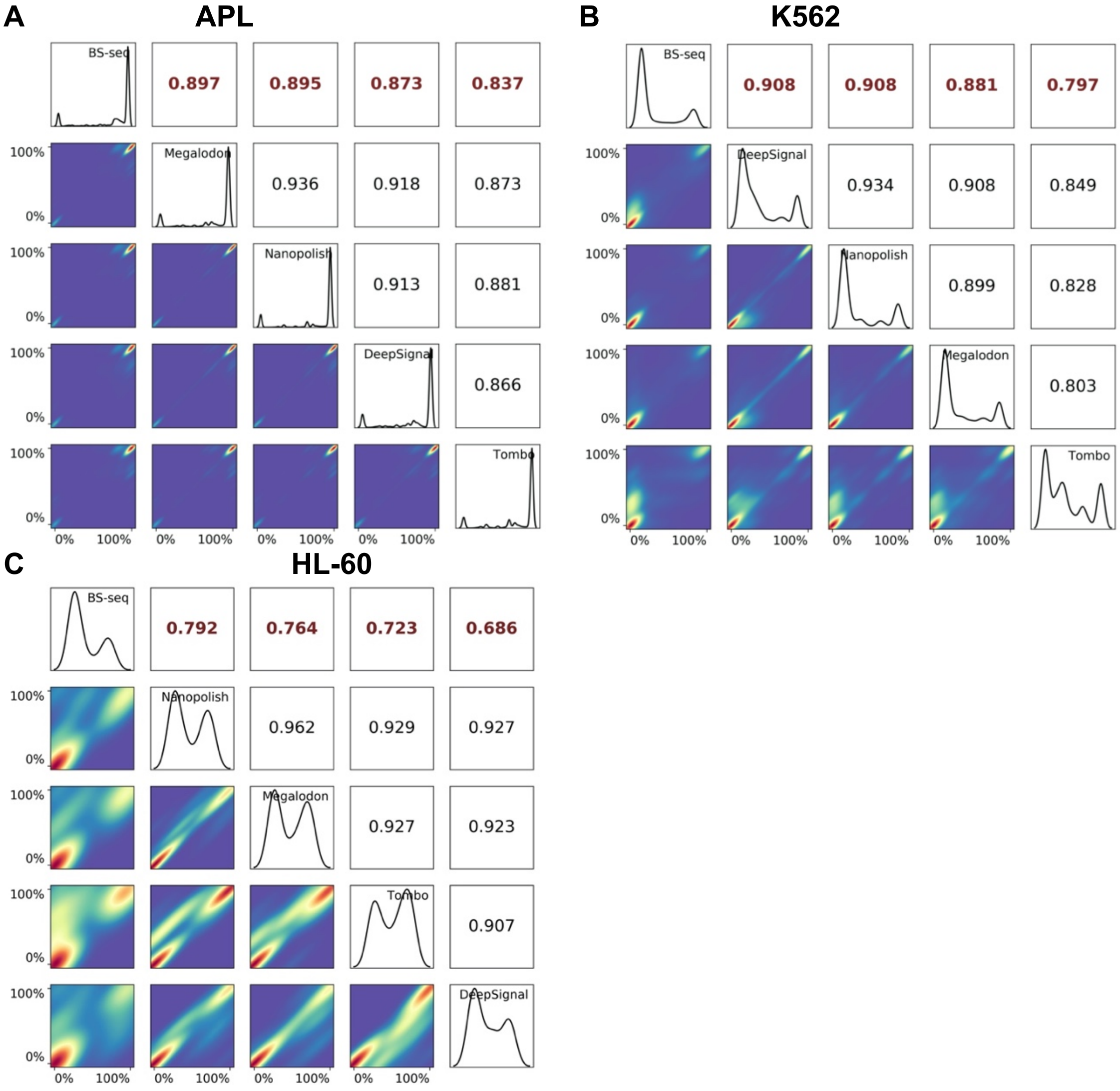


**Figure S5: Correlation plot showing all four tools (read coverage>=3) with BS-Seq for APL (A), K562 (B), and HL-60 (C) Nanopore sequencing datasets (coverage>=5).**

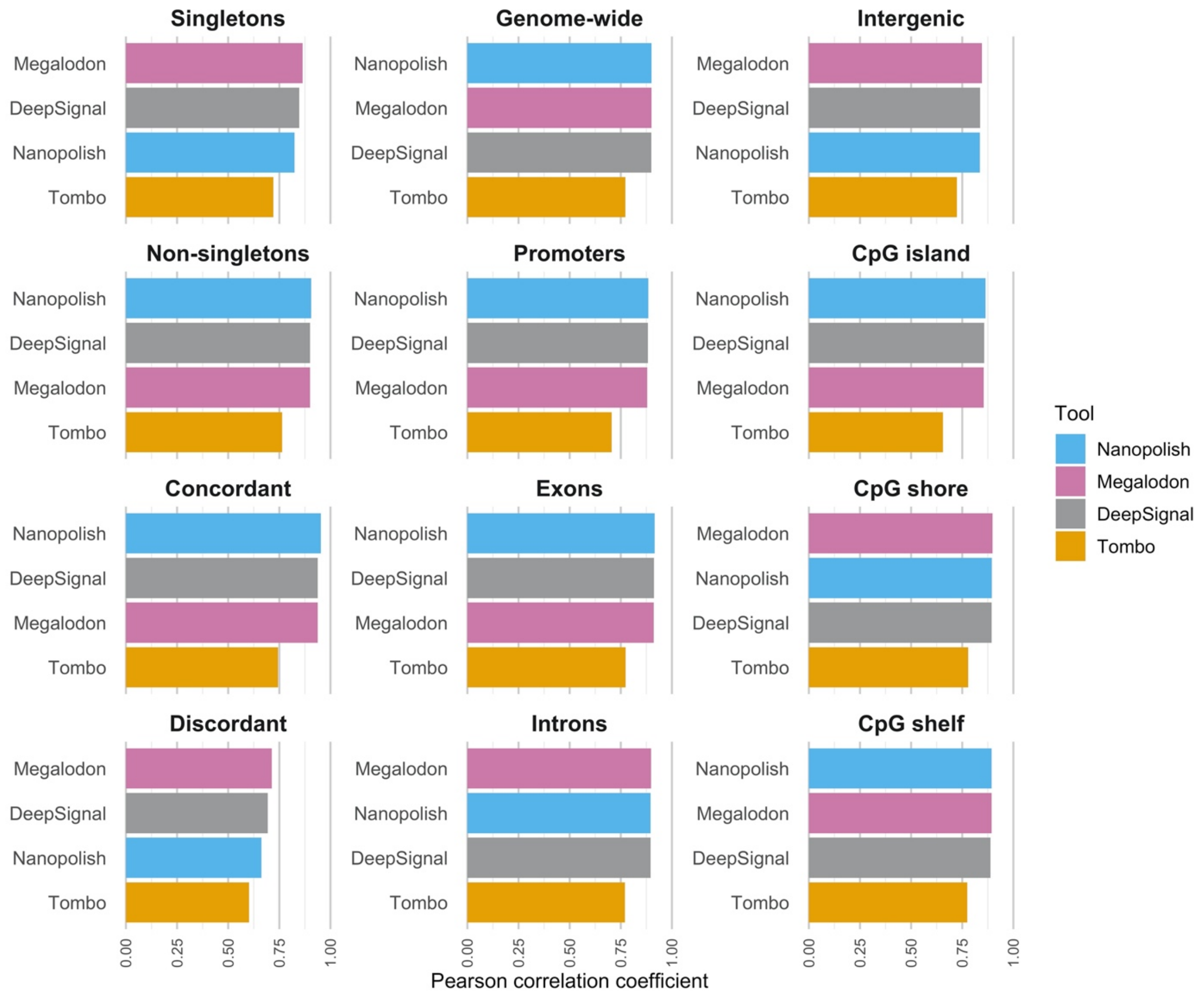


**Figure S6: Correlation comparison across genomic regions for NA19240 dataset.**

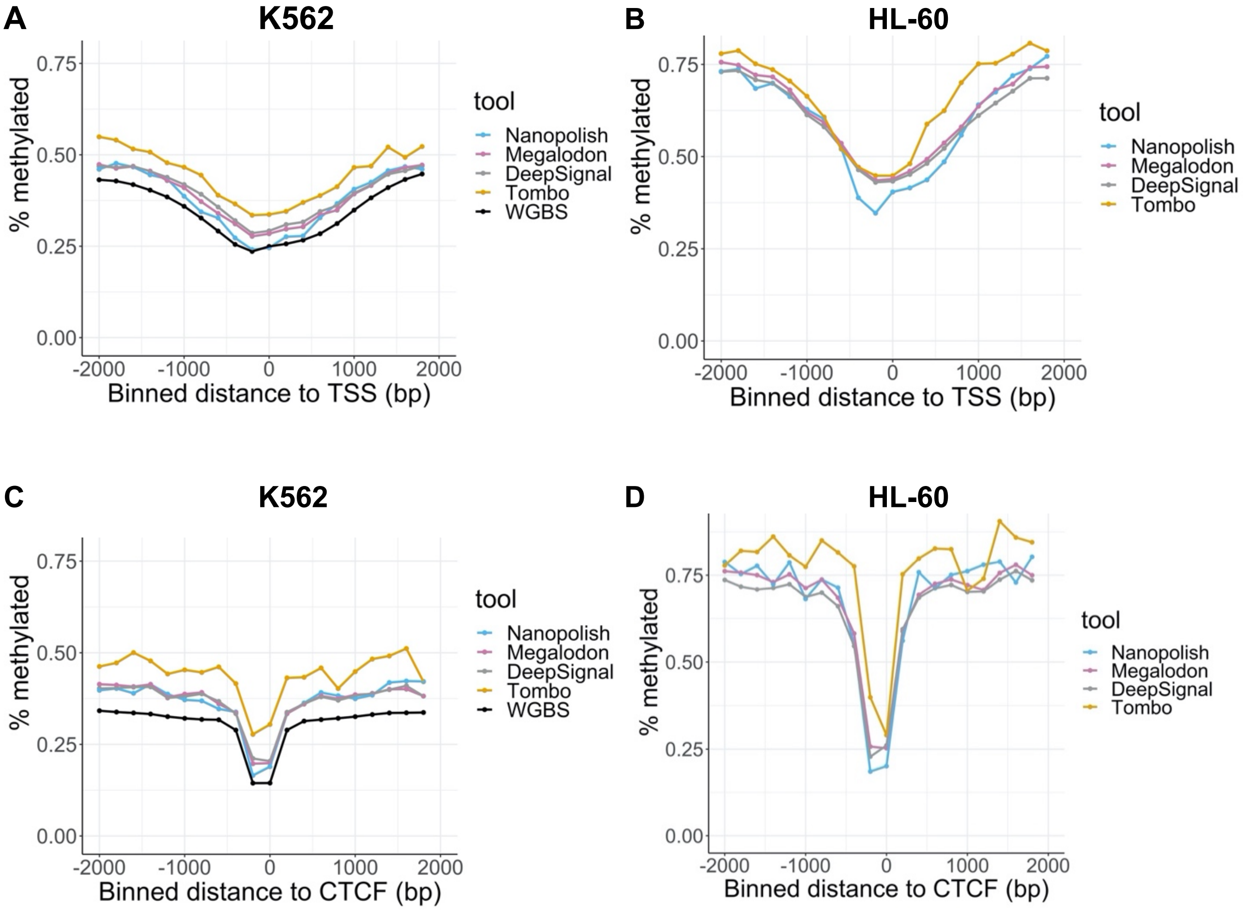


**Figure S7: Relationship between CpG methylation percentage and distance to annotated (A-B) TSS (bin size = 200 bp) and (C-D) CTCF binding peaks in K562 (bin size = 200).**

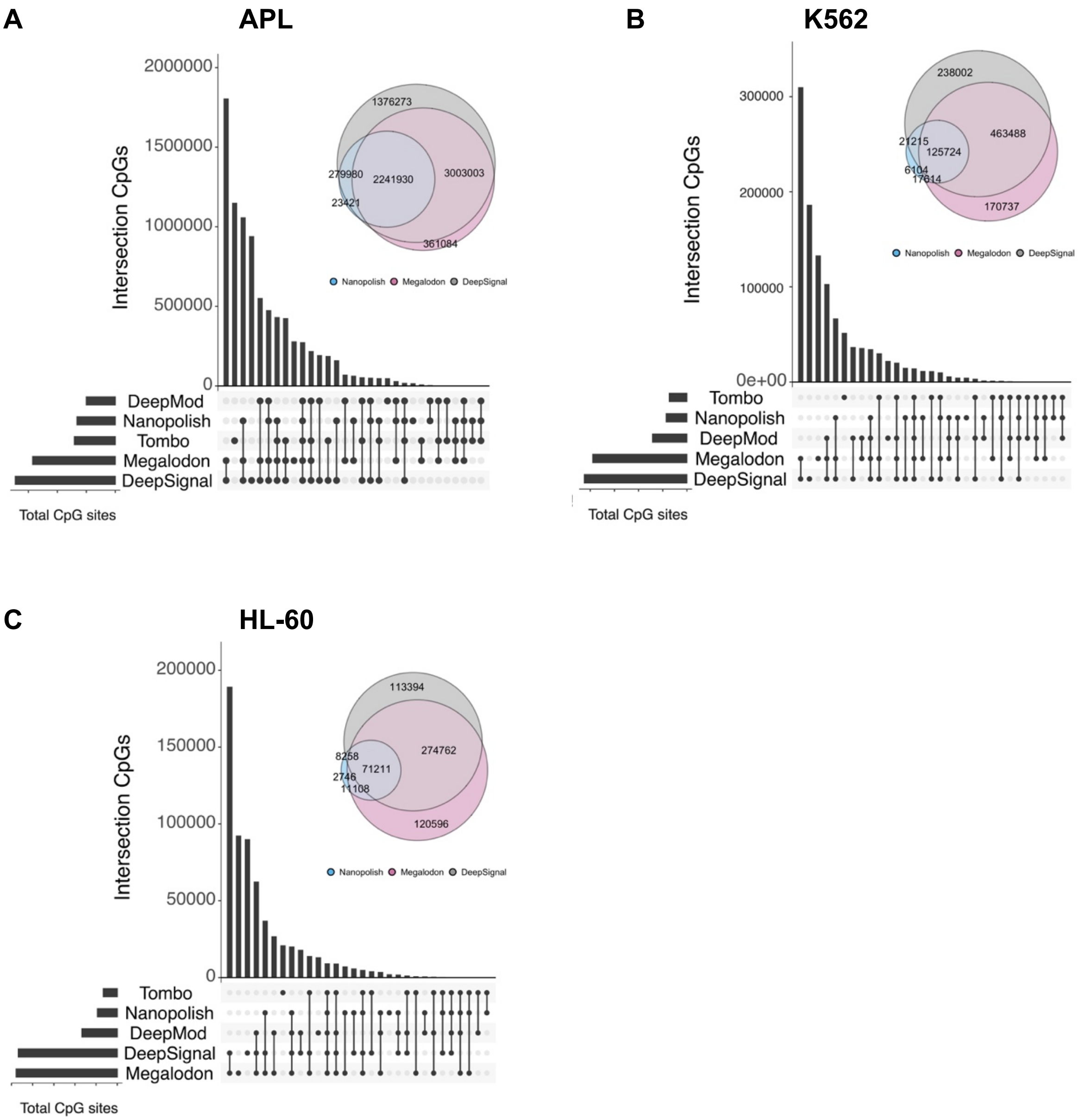


**Figure S8: CpG sites detected by methylation calling tools (read coverage >=3) in APL (A), K562 (B), and HL-60 (C) datasets.**
